# The complete genome of lettuce reveals centromere landscape and empowers breeding

**DOI:** 10.1101/2024.05.14.593296

**Authors:** Ke Wang, Jingyun Jin, Jingxuan Wang, Xinrui Wang, Jie Sun, Dian Meng, Xiangfeng Wang, Li Guo

## Abstract

Lettuce (*Lactuca sativa* L.) is a globally grown vegetable crop as the most popular salad ingredient. Lettuce breeding is crucial to enhancing yield, nutrition and disease resistance, but impeded by lacking a high-quality genome resource. In addition, the released genomes remain fragmented (with hundreds of gaps) and incomplete with omitting pivotal centromeres and telomeres. Here, we report a 2.59 Gb complete telomere-to-telomere (T2T) genome assembly for lettuce. T2T genome analysis revealed the unique genetic and epigenetic architectures of centromere repeats dominated by retrotransposons and satellites. The complete genome and annotation facilitate the identification of candidate disease resistance (NLR) genes induced during grey mold infection. The genomic resources, and biological insights into genetic and epigenetic landscape of centromeres not only advance lettuce research and breeding, but also propel the biology and evolution of centromeres across plant kingdom.

## Introduction

Lettuce (*L. sativa* L.) is an annual plant belonging to Asteraceae family, often used as fresh-cut vegetable and the most popular ingredient in salads. Rich in vitamins, minerals, polyphenols and carotenoids (Yang et al., 2022), it provides numerous health benefits (Shi et al., 2022). In 2021, lettuce has a gross production value of $16.6 billion worldwide, with China, USA and Western Europe as the top three lettuce producers (Food and Agriculture Organization of the United Nations, 2023). Cultivated lettuce is mostly inbred (2n=18) with limited genetic diversity, causing lettuce vulnerable to various abiotic and biotic stresses (Galieni et al., 2015; Richard N., 2004). Hence, lettuce breeding commonly targets improving yield, quality and disease resistance of lettuce, which heavily relies on genetic (*e*.*g*. molecular markers) and genomic (*e*.*g*. reference genome and multi-omics) resources. The first lettuce genome was assembled using next-generation sequencing (NGS) reads in 2017 (Reyes-Chin-Wo et al., 2017). The latest improved lettuce reference genome v11 of the genome for crisphead lettuce cultivar *Salinas* (GCA_002870075.4) was released in 2022, subsequently Shen et al assembled the genome of stem lettuce (*L. sativa var. Augustana*) (Shen et al., 2023). Although these assemblies have greatly facilitated lettuce research (Gao et al., 2022; Pink et al., 2022; Shen et al., 2023; Wei et al., 2021), they remain highly fragmented and incomplete, containing hundreds of gaps and omitting centromeres, rDNA and telomeres, and thus creating a bottleneck to genomic research, gene cloning and molecular breeding.

Given the susceptibility of cultivated lettuce to diseases, the development of disease-resistant cultivars emerges as a critical strategy for environmentally friendly disease control. NLR proteins are widely recognized for their pivotal roles in plant immunity against pathogens (Chou et al., 2023), making them a promising genetic resource for disease control efforts. However, the exploration of NLR genes in lettuce has been hindered by the incompleteness of existing genomes, requiring the advent of a complete lettuce genome.

In this study, we report the first complete T2T genome of the *L. sativa* cv. *PKU06* followed by the identification of centromeres and NLR genes. Our findings revealed that repetitive elements (REs) constitute the majority of the genome and are particularly abundant in the centromeres, with long terminal repeat (LTR) retrotransposons being the predominant type. The 3D genomics of lettuce demonstrated distinct genomic and epigenomic profiles for A/B compartments. In addition, we identified NLR genes including those located within previously unfilled gap regions on lettuce genome. Transcriptomic analysis of these genes during grey mold infection highlighted their involvement in infection resistance. Collectively, our study illuminates the lettuce genome, providing a robust genetic resource for lettuce breeding and disease control efforts.

## Results

The lettuce genome had an estimated size of 2,701 Mb according to *k*-mer analysis (Supplemental Figure 1). In total, we obtained 112.4× PacBio high-fidelity (HiFi) long reads, 42.9× Oxford Nanopore Technology (ONT) ultra-long reads (N50 > 100 kb) and 118.8× Hi-C reads (Supplemental Table 1). Genome assembly was performed using an in-house pipeline (Supplemental Figure 2). First, the HiFi and ONT reads were assembled using hifiasm, resulting in a draft genome consisting 125 contigs. Following decontamination of microbial and plastid sequences, the contigs were anchored on 9 chromosomes using the Hi-C data (Supplemental Figure 3), and error placement or mis-orientation of contigs were manually corrected in Juicebox. This generated a chromosome-scale assembly with only two remaining gaps on Chr4 which were further filled with the ONT reads to generate a gap-free assembly (Supplemental Figure 4). The two nucleolus organizer regions (NORs) on Chr1 and Chr8 were successfully resolved, containing a total of 8.63 Mb rDNA repeat arrays with 884 copies (Figure 1B). The final complete T2T genome (LsT2T) sized 2,593 Mb (Figure 1A) with contig N50 of 283.1 Mb, an 2,264.8% increase in contig N50 compared to 12.5 Mb of *Salinas* (Supplemental Table 2). In addition, we identified all 18 telomeres using the seven-base telomere repeats (CCCTAAA or TTTAGGG) (Supplemental Table 3). LsT2T was highly syntenic (96.96%) to *Salinas* genome albeit with structural variants detected probably due to two different cultivars (Supplemental Figure 5). Importantly, LsT2T closed 384 gaps left in *Salinas* genome and thus substantially improved the contiguity of lettuce genome (Supplemental Table 2).

**Figure 1.**
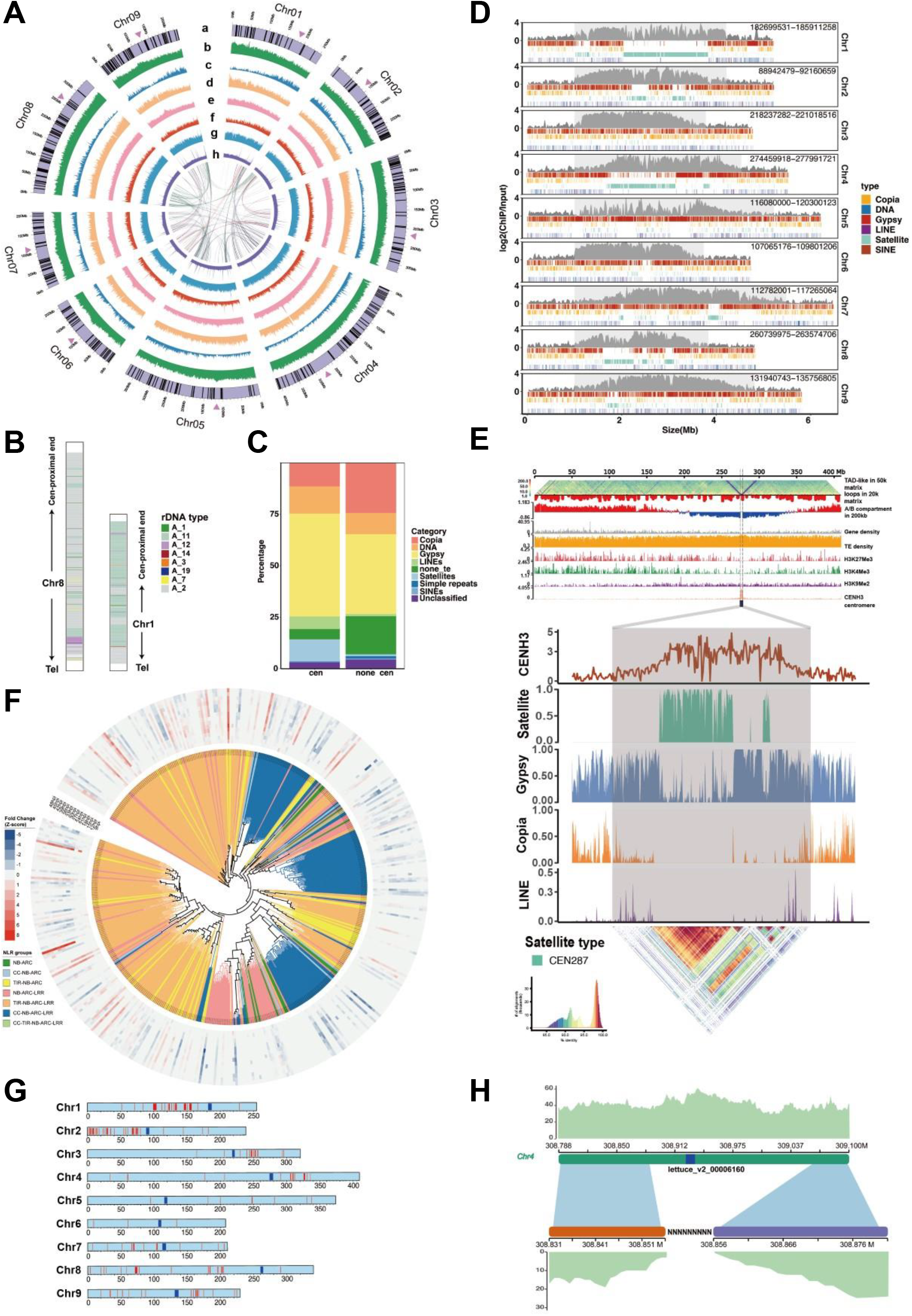
Genemic and epigenomic landscapes of complete telomere-to-telomere genome assembly of lettuce (*Lactuca sativa* L.) (**A**) Concentric circles from outermost to innermost show whole genomic (a) chromosomal feature (arrowheads and black lines represent centromeric location and gaps in *Salinas* genome filled by LsT2T, respectively), (b) GC content density, (c) gene density, (d) LTR-*Copia* density, (e) LTR-*Gypsy* density, (f-h) coverage of HiF, ONT and NGS reads. The innermost color ribbons represent syntenic blocks. (**B**) Graphical summary of the nucleus organizer region (NOR) assemblies. Color blocks represent rDNA variable-length elements types. Tel: telomere. (**C**) Composition of LTRs in centromeric and non-centromeric regions. (**D**) Schematic representation for the distribution of repetitive elements across 9 centromeres. CENH3 ChIP-seq signal (average of two biological replicates) is represented by the enrichment level in 10 kb windows and centromere is encompassed by light grey boxes. From top to bottom track, centromeric LTR-*Gypsy*, LTR-*Copia*, satellite and DNA, LINE, SINE (these three components were merged at bottom track) were marked by red, yellow, gray, blue, purple and brown lines, respectively. (**E**) Genetic and epigenetic profiles (3D genomic feature, gene and TE density histone modification marks) of Chr4 (Other chromosomes are shown in Supplemental Figure 12). From top to bottom of the plot, heatmap of TAD, density of 3D genomic loops and A/B compartments, gene density, TE density, the intensities of H3K27Me3, H3K4Me3, H3K9Me2 histone modifications, ChIP-seq marks and centromeric location were shown. Within the centromere window, ChIP-seq marks, density of satellite, LTR-*Gypsy*, LTR-*Copia*, LINE and sequence identity heatmap were shown from top to bottom. (**F**) Phylogenetic tree for NLR genes constructed using the NB-ARC domain. The NLRs are classified into 7 groups based on domain structures and distinguished by different colors shaded on the tree. Differential gene expression (fold change normalized using z-score) is shown by a heatmap (blue to red), based on a transcriptome dataset of *Botrytis cinerea*-infected lettuce leaves at different stages downloaded from PRJNA1013336. (**G**) Genomic locations of the NLR genes in LsT2T genome. Red and blue lines on each chromosome represent NLR genes and centromeres, respectively. (**H**) Exhibition of a NLR gene identified in filled gap region on Chr4 of LsT2T genome.

We conducted extensive validation of LsT2T as the following. First, we examined Hi-C interaction map of LsT2T, showing no obvious structural assembly errors (Supplemental Figure 3). Secondly, we aligned all raw sequencing data to LsT2T, giving mapping rate of 99.9%, 96.4% and 99.9% for the HiFi, ONT and NGS reads, respectively (Supplemental Table 1). All data exhibited overall uniform genomewide read coverage (Figure 1A), indicating a complete and highly accurate assembly. Interestingly, we observed sporadic instances of elevated coverage in the ONT reads (Figure 1A; Supplemental Table 4) corresponding to chloroplast sequences, suggesting plastid genome integration within the nuclear genome. Further, LsT2T showed a quality value of 58, BUSCO score of 97.6 (Supplemental Table 2), demonstrating high accuracy and completeness of LsT2T.

About 2.1 Gb REs occupying 81.4% of LsT2T were annotated, predominated by transposable elements (TEs) (Figure 1C; Supplemental Table 5). Notably, the majority of TEs were long terminal repeat (LTR) retrotransposons, of which *Gypsy* and *Copia* accounted for 37.84% and 27.23% of LsT2T, respectively. In total, 45,507 protein-coding genes (Supplemental Table 6) were predicted in LsT2T with *ab initio* prediction, homologous proteins and transcriptomic data obtained from five different tissues sequenced by NGS and PacBio Iso-seq. 48.8% of genes were functionally annotated using eggNOG-mapper, and 57.3% were expressed in at least one tissue under the threshold of TPM >= 1 (Supplemental Table 6). We observed that the newly assembled sequences in LsT2T compared to *Salinas* consisted of genes (2.09%), REs (31.34%), centromeres (16.9%) and rDNA arrays (43.4%) (Supplemental Figure 6B), highlighting the importance of a complete genome for unveiling crucial genomic regions.

Centromeres are repeat-rich heterochromatic regions critical to a faithful chromosome segregation during cell division (Cleveland et al., 2003). Centromeres of lettuce were identified by ChIP-seq profiling with a lettuce-specific CENH3 (centromere-specific histone 3) antibody. Lettuce CENH3 ChIP-seq signals well delineated the boundaries of nine centromeres (Figure 1D; Supplemental Table 7) with sizes ranging from 2.7 Mb (Chr6) to 4.5 Mb (Chr7). Centromere position in each chromosome was varied with the ratio of long vs. short ranging from being 1.1 (Chr6) to 3.2 (Chr8) (Figure 1A; Supplemental Figure 5A). Low sequence similarity was observed among the centromeres, suggesting their strong diversification (Supplemental Figure 7). Centromeric repeats were mostly composed of *Gypsy* (56.6%), *Copia* (13.1%) and satellites (16.3%), different from those of non-centromeric regions (Figure 1C). In addition, centromeric *Gypsy* was dominated by *Tekay, Angela* and centromeric retrotransposons of maize (*CRM*) (Supplemental Figure 8A). Notably, *CRM* appeared more frequently in centromeric regions rather in non-centromeric, consistent with previous reports for maize and cotton (Chang et al., 2023; Chen et al., 2023). Phylogenetic analysis of *Gypsy* demonstrated that centromeric *CRM*s formed a unique clade, suggesting centromeric *CRM* expansion without non-centromeric *CRM* (Supplemental Figure 8B). The proportions of satellites in the centromeres varied from 3.25% (Chr3) to 60.14% (Chr1) (Figure 1D; Supplemental Table 7). A *de novo* identification of centromeric satellite monomers with TRASH revealed that 30 bp, 62 bp, 287 bp and 123 bp monomers were predominated satellites (Supplemental Figure 8C). Notably, we observed higher-order repeats (Figure 1E; Supplemental Figure 9) which were mostly composed of 62 bp monomers with miscellaneous short repeats (Supplemental Figure 8C). Analysis of CENH3 enrichment showed that CENH3 preferentially bond to *Gypsy* and satellites (Figure 1E; Supplemental Figures 9 and 10), demonstrating their importance in centromere function.

Despite the decoded lettuce genome, its 3D genomic landscape remains unknown so far. We applied miniMDS for modeling the 3D structure of lettuce genome and using high-resolution Hi-C data (Supplemental Figure 11). The 2.59 Gb lettuce genome was organized into topological associated domains and A/B compartments with low frequency of A/B switching and all centromeres were localized in B compartment (Figure 1E; Supplemental Figure 12). The A compartment had higher gene density and lower TE density than B compartment and two compartments displayed distinctive epigenetic markers (Figure 1E; Supplemental Figure 12). ChIP-seq of histone modifications revealed that H3K4me3 and H3K27me3 marking transcription activation and repression, respectively, were enriched on the A compartments, whereas B compartments were enriched with H3K9me2 typically associated with heterochromatin (Figure 1E; Supplemental Figure 12), a conserved pattern consistent with most plant 3D genomes reported so far.

We systematically identified 514 putative NLR genes in LsT2T and classified them into seven subfamilies via phylogenetic analysis of NB-ARC domain (Figure 1F), whereas the same approach identified only 484 NLR genes in *Salinas* genome. The majority of LsT2T NLR genes were tandemly duplicated and genomically clustered, particularly on Chr1 and Chr2 (Figure 1G). Interestingly, four new NLRs were found in the filled gap regions of LsT2T (Figure 1H; Supplemental Figure 13), including one localized within a gap region of Chr4 where only LsT2T had ONT read coverage. Transcriptomic analysis revealed that 58 NLRs were significantly upregulated during grey mold (*Botrytis cineara*) infection compared to the mock, in which 38 NLRs encoding TIR-NB-ARC(-LRR) domains were predominant (Figure 1F; Supplemental Table 8). The most significantly upregulated NLR lettuce_v2_00029769 was homologous to *Arabidopsis thaliana* AT5G36930 encoding a TIR-NB-ARC-LRR type NLR. Future functional characterization of the infection-induced NLR genes revealed by T2T genome will provide insight into lettuce immunity against pathogens.

In summary, we generated the complete T2T genome of lettuce, the first for Asterids, and dissected the complex genetic and epigenetic landscape of centromeres, which will serve as an essential resource for expediting lettuce research and genetic improvement.

## Materials and Methods

### Plant material and growth conditions

*L. sativa* plants were grown in regular azalea pots filled with a combination of potting mix, clay and vermiculite in greenhouse of Peking University Institute of Advanced Agricultural Sciences (Weifang, China). The fresh leaves of seven weeks old *L. sativa* were harvested and used for DNA extraction and sequencing. Tissues of leaf, root, stem, and flower at two days post anthesis, and seeds at 21 days post anthesis for *L. sativa* were collected for RNA extraction and sequencing.

### Extraction of DNA and RNA

The extraction of high molecular weight (HMW) genomic DNA was performed using CTAB method (Z. Li et al., 2020). The isolation of total RNA was performed using TRIzol™ Reagent (Thermo Fisher Scientific, USA) following manufacturer’s instruction. RNA was assessed using RNA Nano 6000 Assay Kit (Agilent Technologies, USA) following manufacturer’s instruction.

### Genome and transcriptome sequencing

For short read sequencing, 5 μg of HMW DNA sample was fragmented by sonication to a size of 350 bp and sequencing library was prepared using NEB Next® Ultra™ DNA Library Prep Kit for Illumina (New England Biolabs, USA) following manufacturer’s instruction, and sequenced using Illumina Novoseq 6000 platform. For PacBio HiFi sequencing, 15 μg of HMW DNA was sheared by g-TUBE (Covaris, USA) and applied for PacBio SMRTbell library. The library was size-selected with 15 kb as a cutoff and sequenced using PacBio Sequel II system. For Oxford Nanopore sequencing, 1.5 μg of HMW DNA was size-selected with 3 kb as a cutoff, and applied for Ligation Sequencing Kit (SQK-LSK109) (Oxford Nanopore Technologies, UK). The final DNA library was sequenced using the GridION X5/PromethION sequencer (Oxford Nanopore Technologies, UK) by Single-Molecule Sequencing Platform at Peking University Institute of Advanced Agricultural Sciences (Weifang, China). The preparation of Hi-C library was performed following the standard protocol described previously with some modifications (Belton et al., 2012). The library was sequenced using Illumina PE150 (Illumina, USA) to obtain 2×150 bp paired-end reads. For transcriptome sequencing library construction, a total of 3 μg of RNA was applied for VAHTS Universal V6 RNA-seq Library Prep Kit for Illumina (Vazyme Biotech, China) following the manufacturer’s instruction. The library was sequenced using HiSeq X Ten (Illumina, USA) with paired-end sequencing program (PE150).

### Genome assembly

After counting k-mers with Jellyfish (Marçais & Kingsford, 2011) (v2.3.0), the genome size of *L. sativa* was estimated by k-mer frequency analysis with GenomeScope (Vurture et al., 2017) (v1.0). Multiple strategies deploying different algorithms were used for the *de novo* assembly of the genome. Briefly, the HiFi reads were subjected to hifiasm (Cheng et al., 2021) (v0.16.1) to generate a draft assembly (L1). Both the HiFi reads and the ONT reads were self-corrected, trimmed, and assembled using hifiasm to generate contigs with high quality and long continuity. Then we aligned the contigs to the reference genomes of *L*.*sativa* chloroplast and mitochondria with minimap2 (H. Li et al., 2009a)(v2.24) (-x asm5), removing chloroplast or mitochondria genome sequences from the contigs with at least 50% coverage to generate the cleaned assembly (L2).

To anchor the contigs of L2 assembly, the Hi-C reads were clustered, ordered and oriented using Juicer (Durand, Shamim, et al., 2016) (v1.5) with subsequent manual adjustment using Juicebox (Durand, Robinson, et al., 2016) (v1.11.08) and 3D-DNA (Dudchenko et al., n.d.) (v180419), anchoring the contigs into nine pseudo-chromosomes without any large and obvious assembly errors. The three-dimension structure of lettuce genome was modeled using miniMDS (Rieber & Mahony, 2017) from the Hi-C data.

Given that chloroplast sequence insertions were detected flanking Gap1 (Supplemental Figure 4), we extracted the 100 kb sequence on each side of chloroplast sequence insertion away from the gap, and searched for ONT reads larger than 100 kb that span Gap1. This was done on both ends, and then we looked for intersections in the two ONT read sets, finding ONT reads that can span gap1 to fill Gap1. Gap2 region (Supplemental Figure 4) is a repeating segment of GAA that is not highly conserved. We also took the 100 kb sequences at the left and right ends of the gap to search for potential reads spanning Gap2. As a result, there were no reads that spanned Gap2. Then we took the reads with the longest extension at both ends and found that an intersection. Finally, we used these two reads to fill Gap2.

To polish genome assembly, the HiFi and ONT reads were aligned to the assembly using Winnowmap (Jain et al., 2020), retaining only the primary alignments. Small indels (< 50 bp) were called from the alignment results of HiFi using DeepVariant (Poplin et al., 2018). Structural variations (> 50 bp) were identified from the alignment results of HiFi and ONT using Sniffles (Sedlazeck et al., 2018). Then, Merfin (Formenti et al., 2022) was applied to evaluate the variant calls and BCFtools (https://samtools.github.io/bcftools/) was used to correct the bona fide variants in the genome assemblies.

### Genome validation and quality evaluation

The quality of LsT2T genome was evaluated using following methods. QV was assessed using Merqury (Rhie et al., 2020)(v1.3) (parameters: k = 21 count) and BUSCO (v5.4.7) was evaluated using embryophyta_odb10 database (Manni et al., 2021) (n = 2,326). For mapping statistics, the NGS short reads were mapped using BWA (H. Li & Durbin, 2009)(v0.7.17), and the HiFi and ONT long reads were mapped using minimap2. Samtools (H. Li et al., 2009b) (v1.10) was used to count the mapping rates and coverage depth. All gap regions which were resolved in T2T genome were manually checked in IGV (Robinson et al., 2011) (v2.12.3) windows.

### NOR assembly

Complete rDNA fragments were obtained from LsT2T assembly as a reference to call variants using a modified method described previously (Wlodzimierz et al., 2023). Deploying BLAST alignment, the HiFi reads mapped to the rDNA fragments were selected with 9 kb as a cutoff, acquiring a 900 Mb read set. The read set was annotated using barrnap (https://github.com/tseemann/barrnap), obtaining 5.8S, 18S and 25S rDNA structural information. We uniformly spliced reads into segments at 25S rDNA sequences, mapping to the reference rDNA sequence using minimap2. Next, we manually examined the number of variant types in the alignments, extracting the sequence of the regions where the variant types were located and counted. Then, we aligned the different types of rDNA fragments collected back to the contig assembled by hifiasm containing rDNA tandem repeats through minimap. Two types of rDNA repeats were found at the centromeric end of the NORs on Chr1 and Chr8, respectively. Therefore, we estimated the length of NORs from ONT and HiFi reads respectively, and then divided the remaining hifiasm contigs into two categories according to the rDNA type and assigned them to the corresponding NOR regions.

### RNA-seq data analysis

The transcriptome data were mapped to the LsT2T genome using HISAT2 (Kim et al., 2019) (v2.1.0) (default parameters). The number of reads on each gene was counted by HTSeq (Anders et al., 2015) (v0.12.3). The gene expression levels in all samples were calculated using Transcripts Per Kilobase of exon model per Million mapped reads (TPM). Differentially expressed genes were analyzed using DESeq2 (Love et al., 2014) package in R.

### Genome annotation

Protein-coding genes were predicted using MAKER (Cantarel et al., 2008) (v2.31.11) pipeline with a combination of ab initio prediction, homologous protein, RNA-seq evidences in two successive rounds. In the first round, based on given RNA-seq data and homology proteins, MAKER deployed BLAST to align transcripts and proteins to the soft-masked genome, using Exonerate (Slater & Birney, 2005) (v2.2.0) to polish the BLAST hits and accurately annotate the coding regions. A subset of MAKER gene models with AED scores < 0.25 was used to train SNAP (Korf, 2004) for three rounds. GeneMark-ET and Augustus models were trained using BRAKER (Hoff et al., 2019) (v2.1.6) pipeline. To implement MAKER and BRAKER pipelines, the RNA-seq data were assembled into a transcriptome using Trinity (Grabherr et al., 2011)(v2.8.4) and were aligned to genome assembly through HISAT2 (v2.1.0). The PacBio long-read transcriptome data were processed using the SMRT Analysis software Isoseq3 (https://github.com/PacificBiosciences/IsoSeq).

### REs analysis

LsT2T assembly was applied for REs annotation using Repbase database (Bao et al., 2015), constructing a *L. sativa* specific *de novo* repeat library using RepeatModeler (Flynn et al., 2020). Repetitive elements in the genome were annotated and masked by a combination of TEsorter (Zhang et al., 2022), DeepTE (Yan et al., 2020), RepeatMasker (Tarailo-Graovac & Chen, 2009) (v4.1.2) (parameters: -xsmall -s -no_id -cutoff 255 -frag 20000 -e ncbi). LTR_Finder (Xu & Wang, 2007) (v1.2), LTRharvest (Ellinghaus et al., 2008) (v1.6.2), and LTR_retriever (Ou & Jiang, 2018) (v2.9.0) were used to identify LTR elements.

### CENH3 ChIP-seq and data analysis

Leaves of lettuce were fixed with 1% formaldehyde solution in MS buffer (10 mM potassium phosphate (pH 7.0), 50 mM NaCl), followed by incubation with 0.15 M glycine under vacuum. The ChIP experiment was performed following the previously described method (Guo et al., 2024) using lettuce anti-CENH3 antibody prepared in rabbit. The ChIP-seq libraries were sequenced using Illumina Novaseq 6000 platform. The ChIP-seq data were analyzed as previously described (Guo et al., 2024).

### Genomewide identification of NLR genes

NLR-Annotator (Steuernagel et al., 2020) (v.2.1b) was deployed to identify genomic segments containing putative nucleotide-binding domain and leucine-rich repeat of NLR genes. The outputs were integrated to the whole-genome annotation with deleting redundant genes, generating a subset NLR genes of lettuce. The sequences of NB-ARC motif were applied for the construction of phylogenetic tree using NLR-Annotator.

## Funding

This project was supported by the Key R&D Program of Shandong Province (ZR202211070163) and the Natural Science Foundation for Distinguished Young Scholars of Shandong Province (ZR2023JQ010). LG is also supported by Taishan Scholars Program of Shandong Province.

## Author Contributions

L.G. conceived and supervised the study. X.W. and D.M. prepared the plant samples and conducted experiment. K.W., J.W., J.S., X.W. conducted bioinformatic analysis. J.J., K.W., J.W. and L.G. wrote and revised the manuscript. All authors have read and approved the manuscript.

## Acknowledgments

We would like to thank the Single-Molecule Sequencing Platform and Bioinformatics Platform at Peking University Institute of Advanced Agricultural Sciences for providing the ONT sequencing and high-performance computing resources.

## Notes

### Competing Interest Statement

The authors have declared no competing interest.

