## Supplemental Figure 1-13 for "The complete genome of lettuce reveals centromere landscape and empowers breeding"

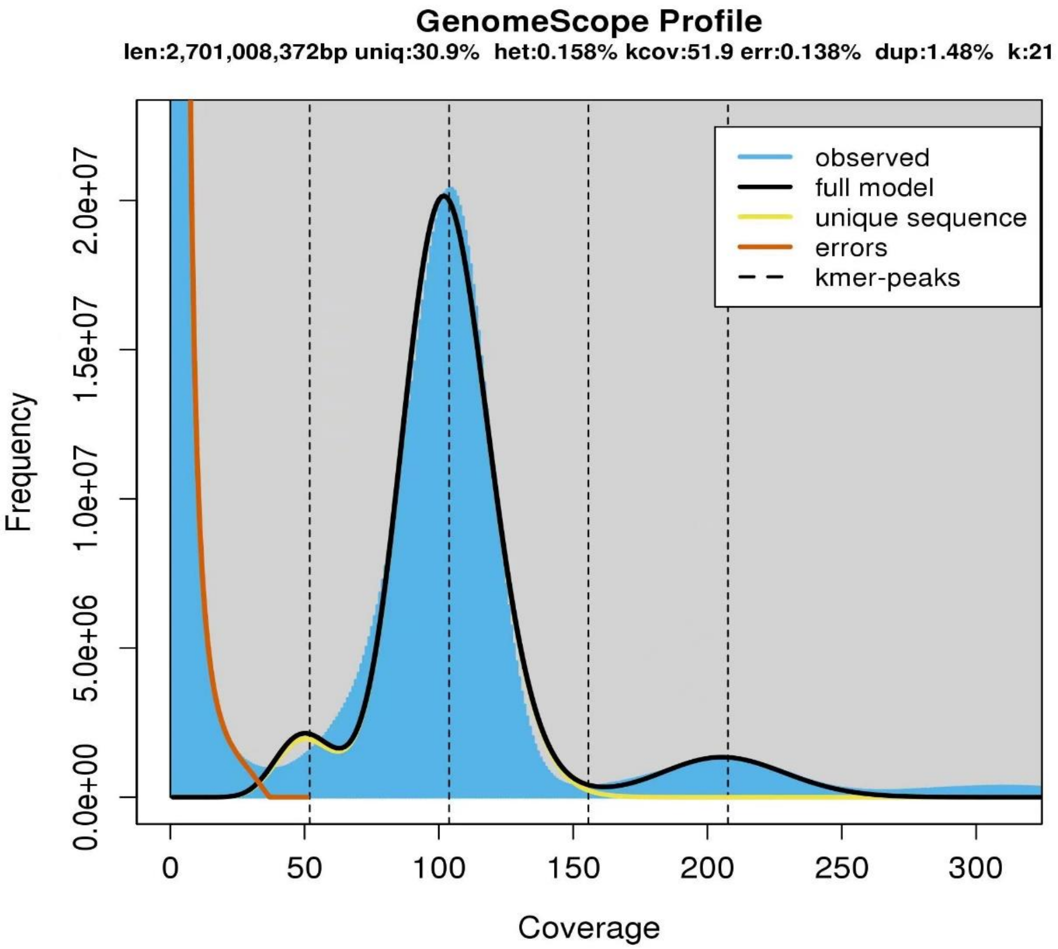


**Supplemental Figure 1 Genome survey of lettuce (*Lactuca sativa* cv. *PKU06*).** kmer (k=21) frequency analysis was performed using Illumina short reads (100x coverage) with Jellyfish and GenomeScope2.

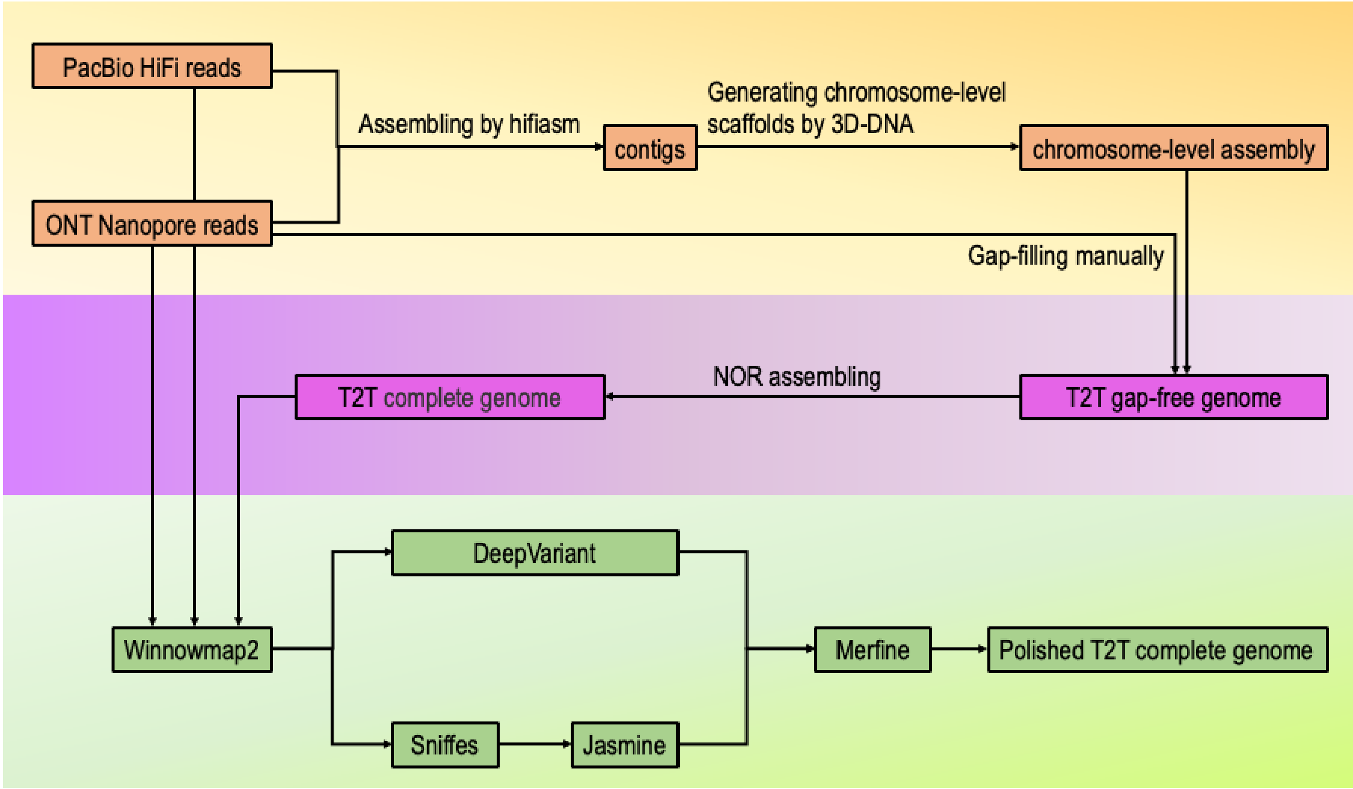


**Supplemental Figure 2. Diagram of genome assembly pipeline used in this study.**

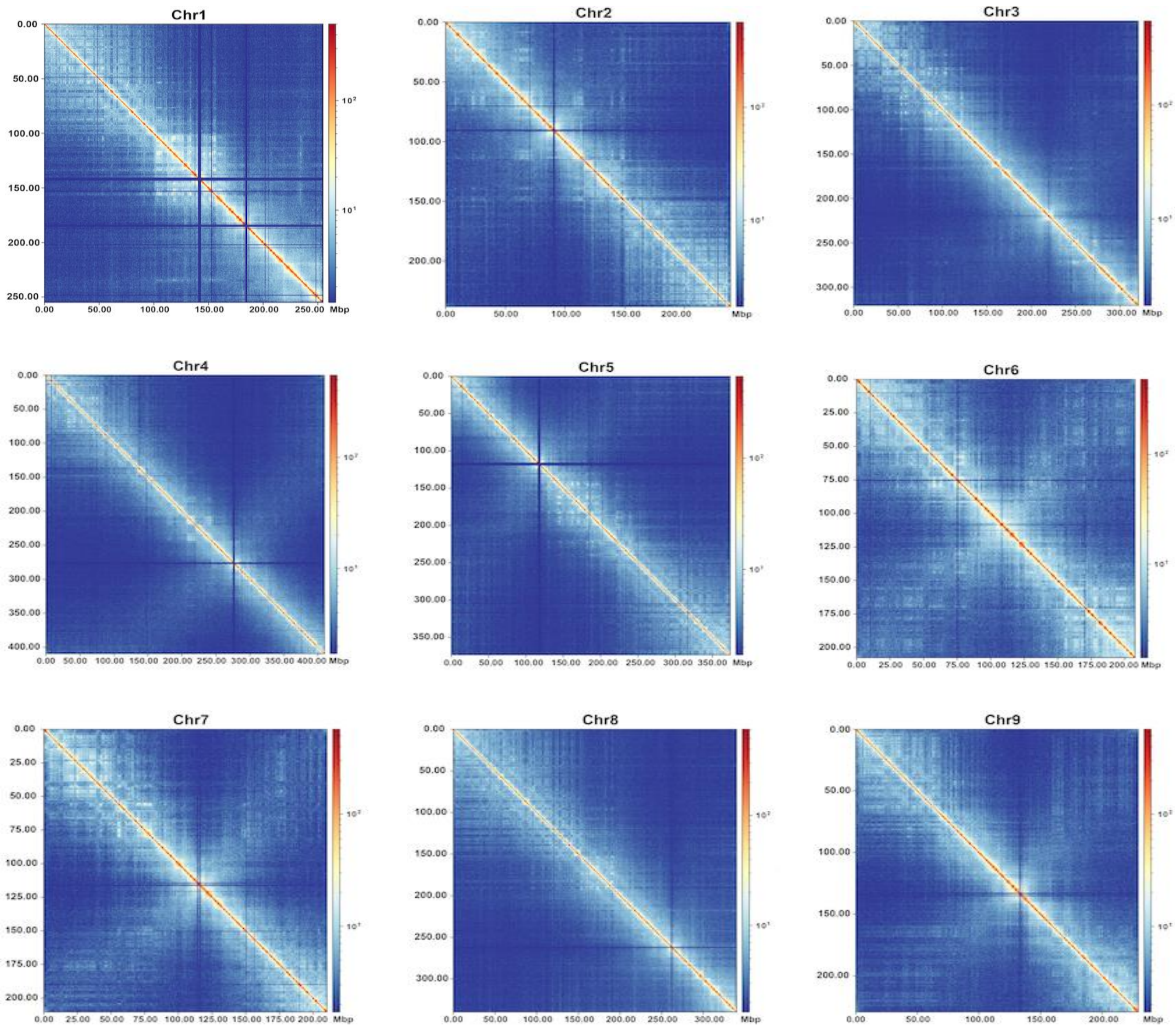


**Supplemental Figure 3. Hi-C interaction heatmaps of LsT2T genome.** The color bars are scaled according to the interaction intensity for each 200-kb bin.

### Gap1

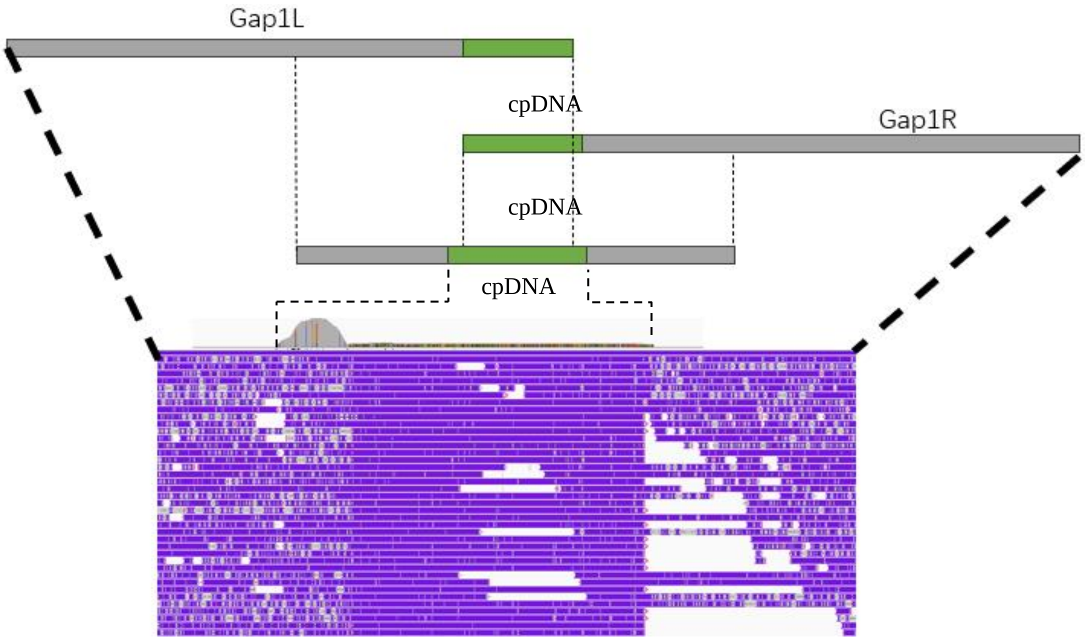


### Gap2

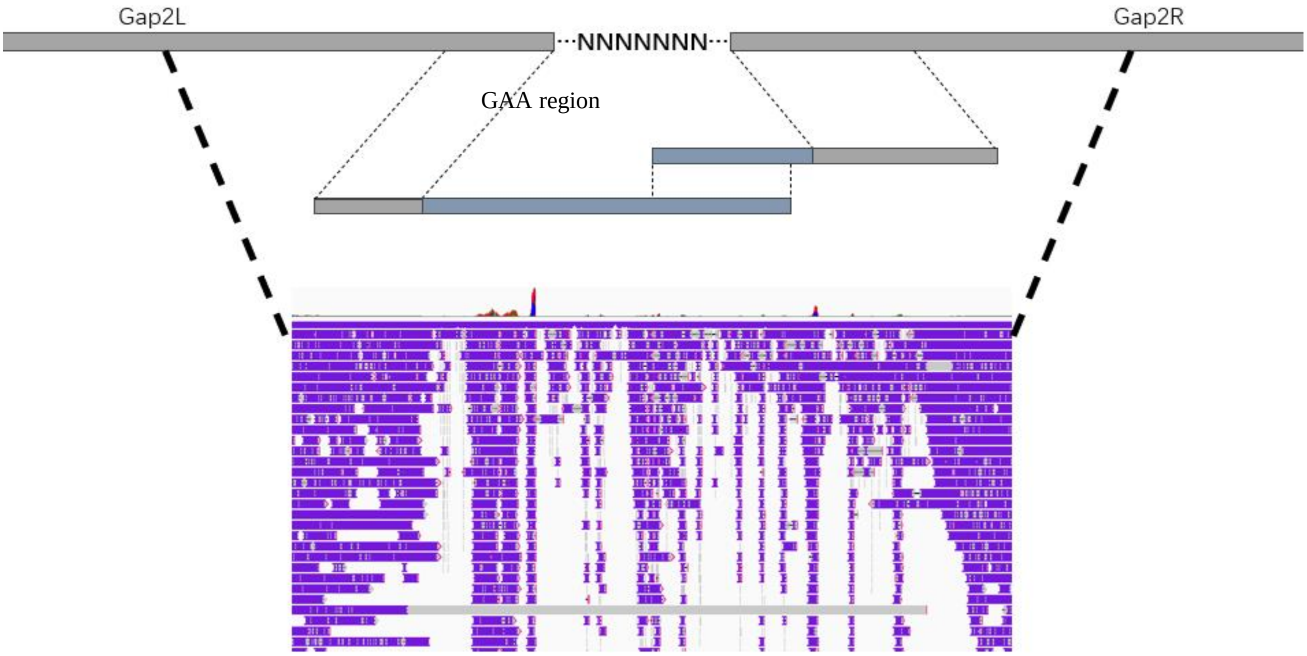


**Supplemental Figure 4. Gap filling diagram for two remaining gaps of initial assembly.** ONT reads spanning the gap regions are utilized for gap filling, and their presences are illustrated in a screenshot of the Integrative Genomics Viewer. cpDNA: chloroplast DNA insertion.

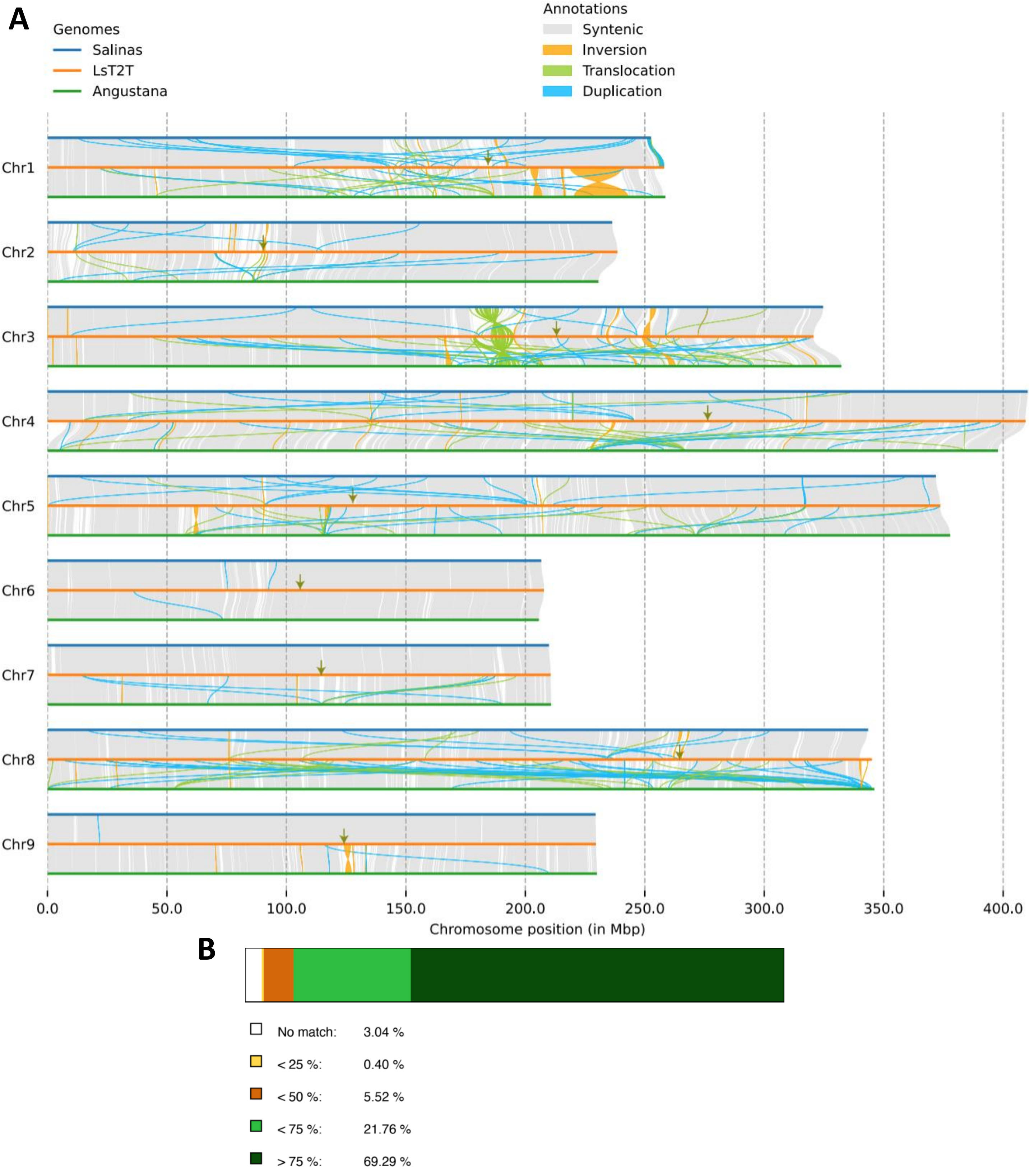


**Supplemental Figure 5. Comparison of genomic feature among LsT2T, *Salinas* and *Angustana* genome assemblies of *L. sativa*.**

(**A**) Syri plot of the genomes showing the syntenies and inversions. The locations of centromere on LsT2T genome were indicated by green arrows.

(**B**) The composition of genome sequence similarity between LsT2T and *Salinas* genomes. The colors represent different level of sequence similarities.

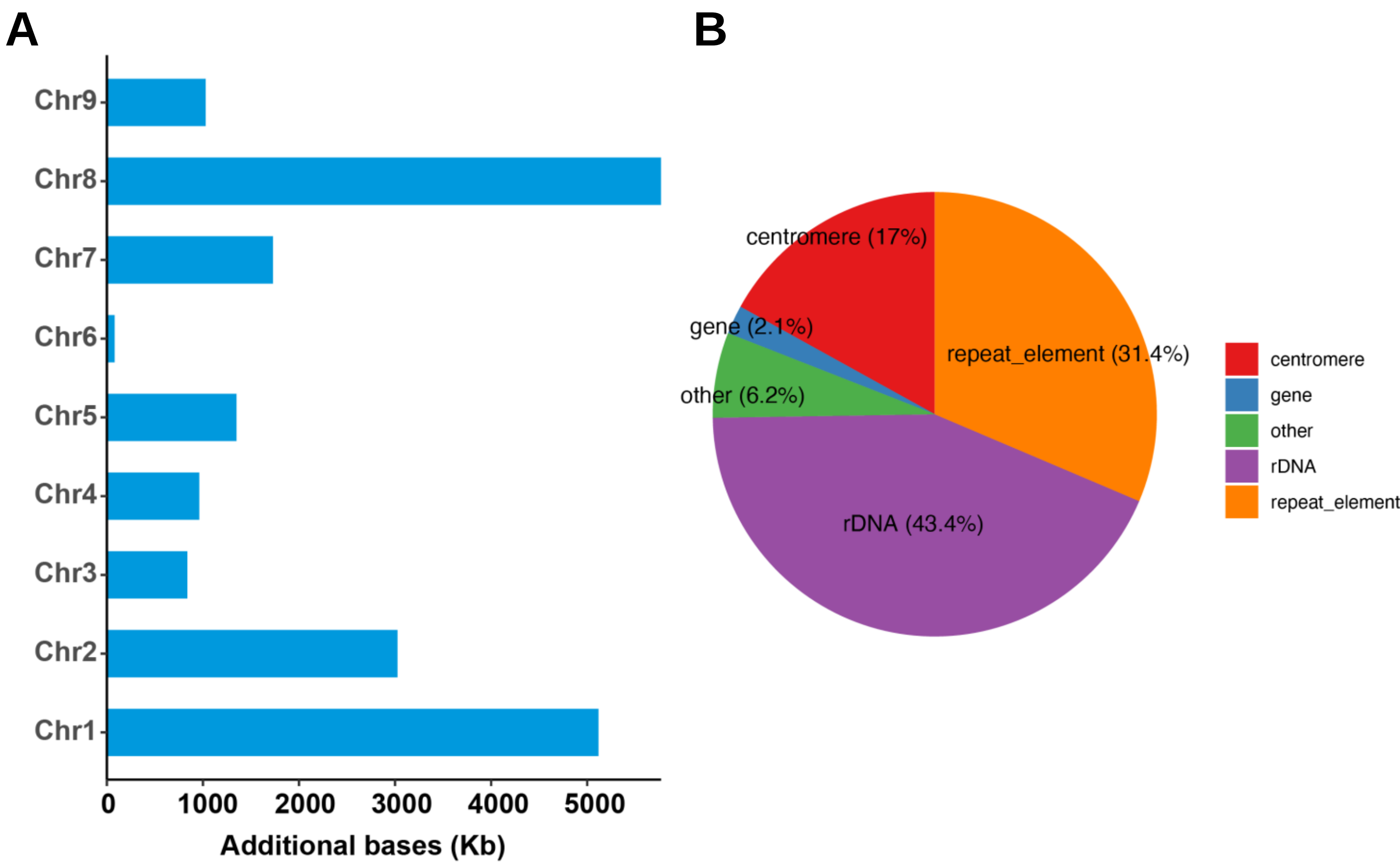


**Supplemental Figure 6. Characterization of newly assembled sequence regions in LsT2T genome.**

(**A**) Statistics on the length of the newly assembled sequence regions by each chromosome.

(**B**) Statistics on the composition of the newly assembled sequence regions.

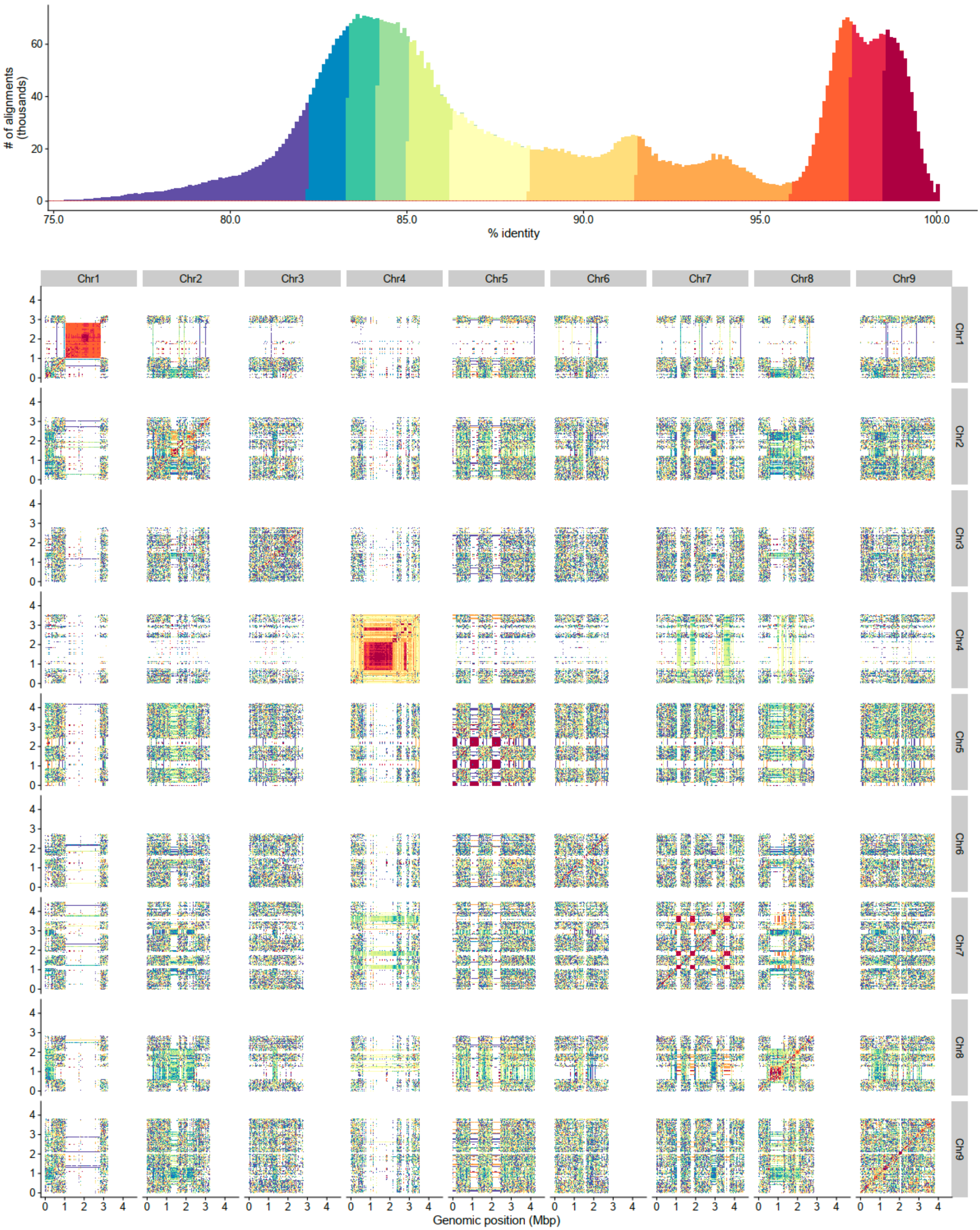


**Supplemental Figure 7. Sequence similarity heatmaps of centromeres across chromosomes in LsT2T genome.** The sequence similarities among the centromeres were visualized using StainedGlass. The heatmap depicts the number of alignments (Y-axis) with sequence similarity (X-axis, 0-100%) is shown at the top.

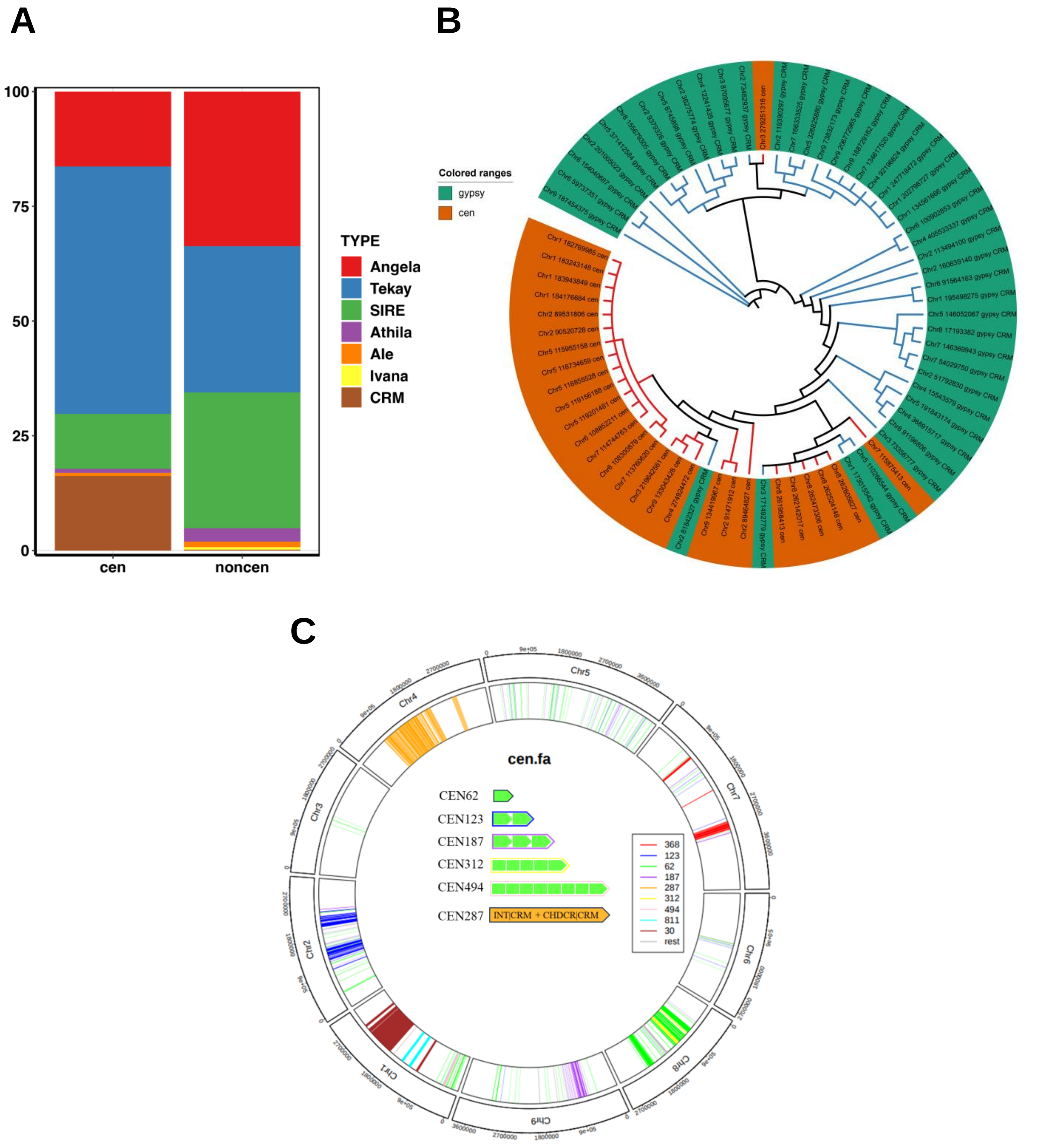


**Supplemental Figure 8. Characterization of REs in LsT2T genome.**

(**A**) LTR composition of non-centromeric and centromeric regions. The colors represent different types of LTR.

(**B**) Phylogenetic tree of *Gypsy* with intact *CRM* domain in non-centromeric and centromeric regions. Emerald and orange represent those of non-centromeric and centromeric regions, respectively.

(**C**) Characterization of sateliite types in  LsT2T genome. The colors represent different types of satellite.

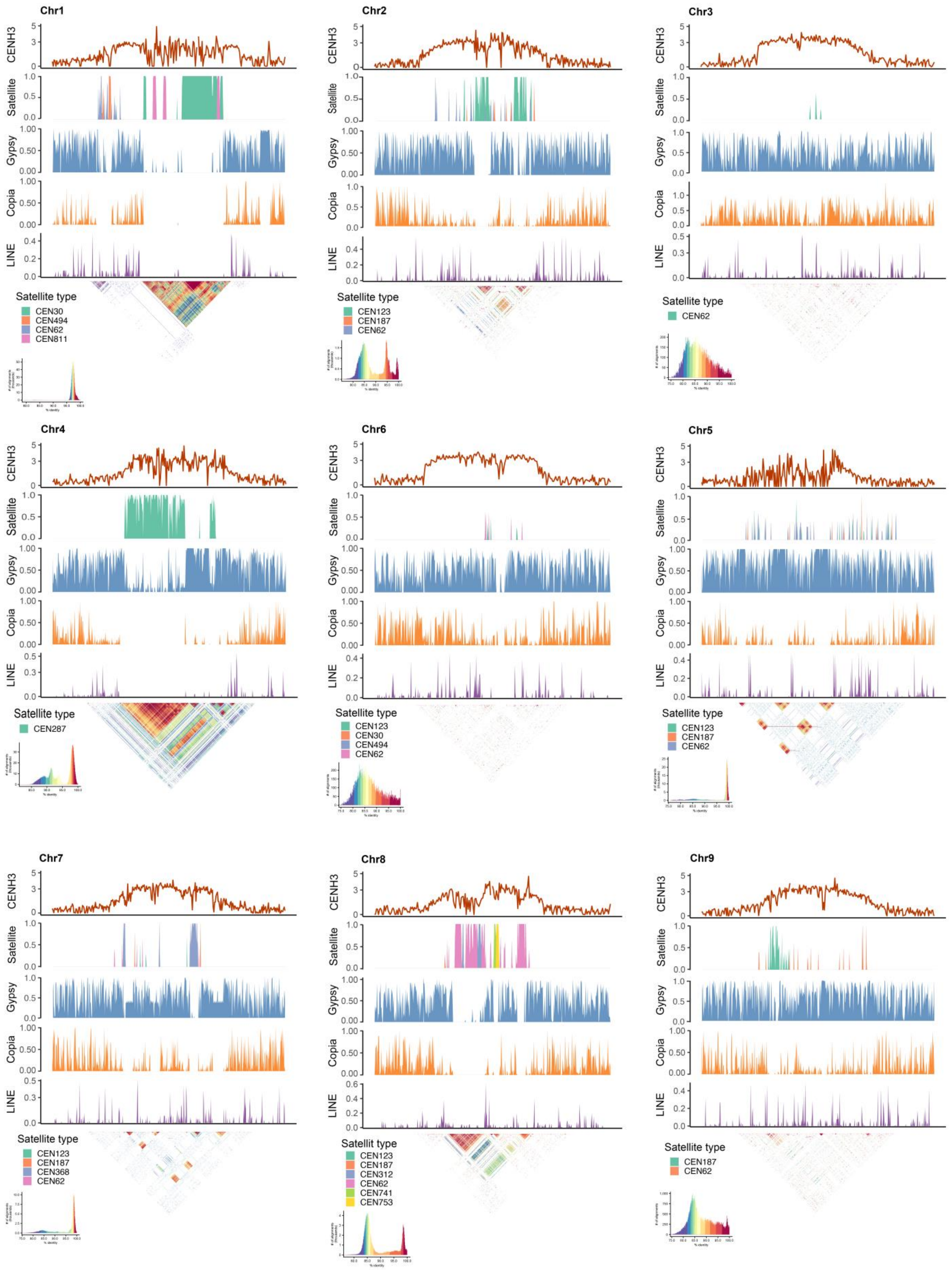


**Supplemental Figure 9. Characterization of centromeric REs composition in LsT2T genome.** The plots for each chromosome showing CENH3 ChIP-seq peaks, the density of satellites (the colors represent different monomers), *Gypsy*, *Copia*, LINE, and sequence identity heatmap which are arranged from top to bottom.

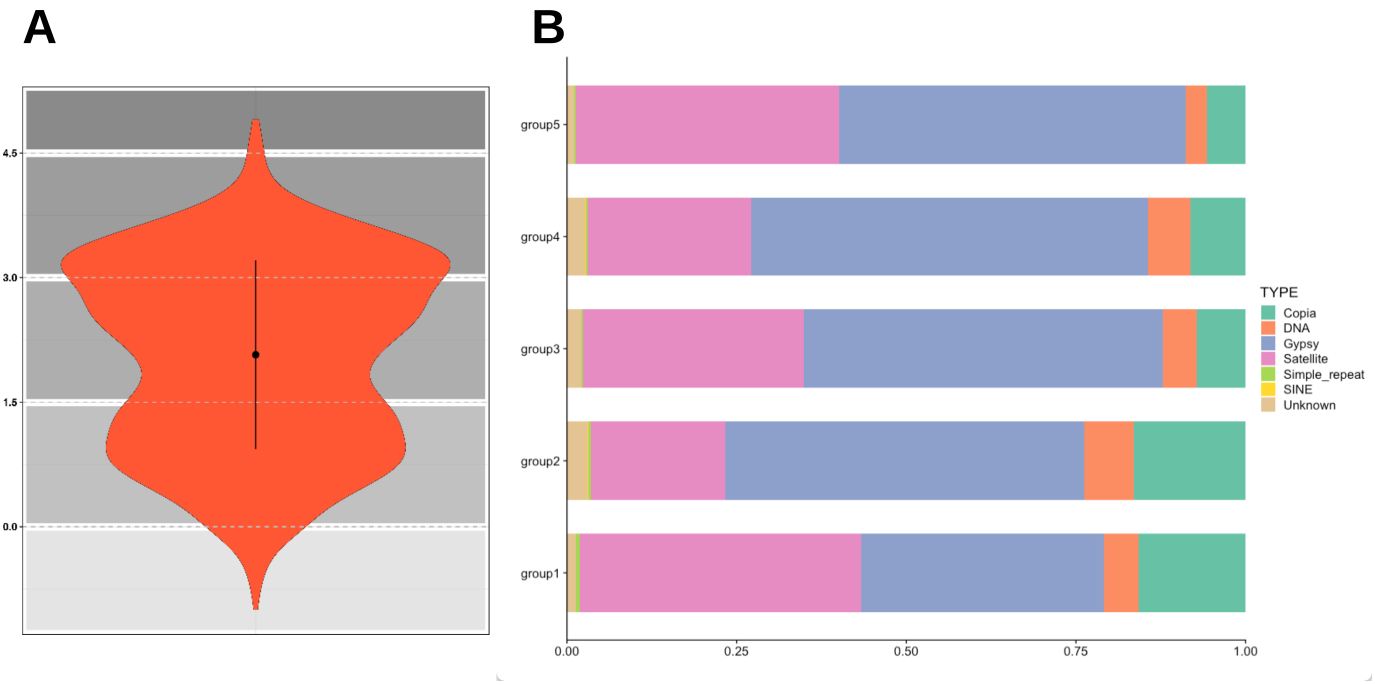


**Supplemental Figure 10. Analysis of CENH3-binding preference for centromeric components.**

(**A**) The profile of CENH3-binding intensities. The distribution of different level intensities were plotted to Y-axis according to log2 value of the ChIP peaks which were normalized against input.

(**B**) Centromeric composition of CENH3-binding regions. Five distinctive groups were categorized based on th intensities: minimum to 0 (group1), 0-1.5 (group2), 1.5-3 (group3), 3-4.5 (group4), 4.5-maximum (group5). The colors indicate the types of centromeric component.

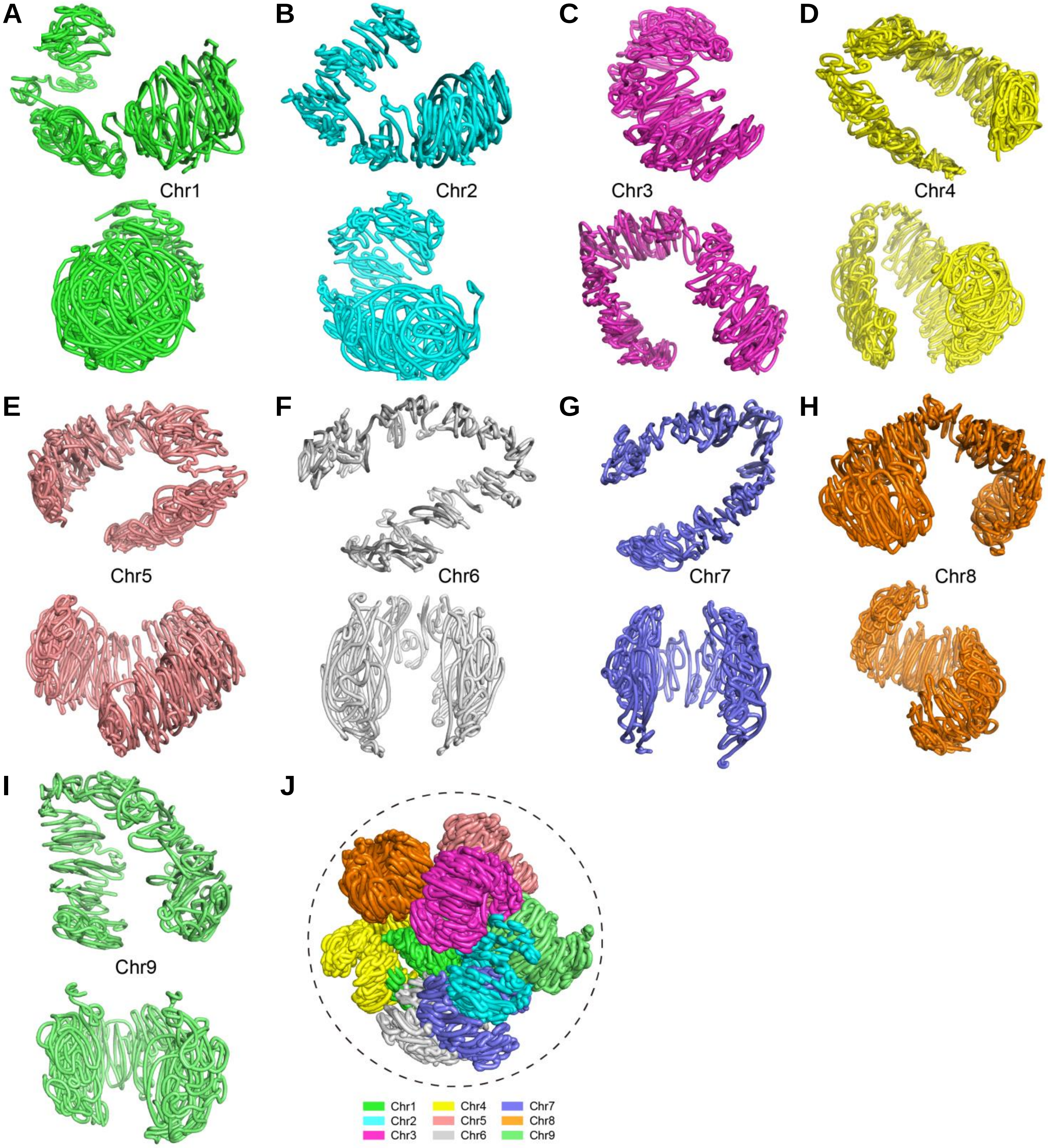


**Supplemental Figure 11. 3D modeling of LsT2T chromosomes.**

(**A-J**) The 3D models of Chr1-9 and whole genome were plotted using iminiMDS, respectively.

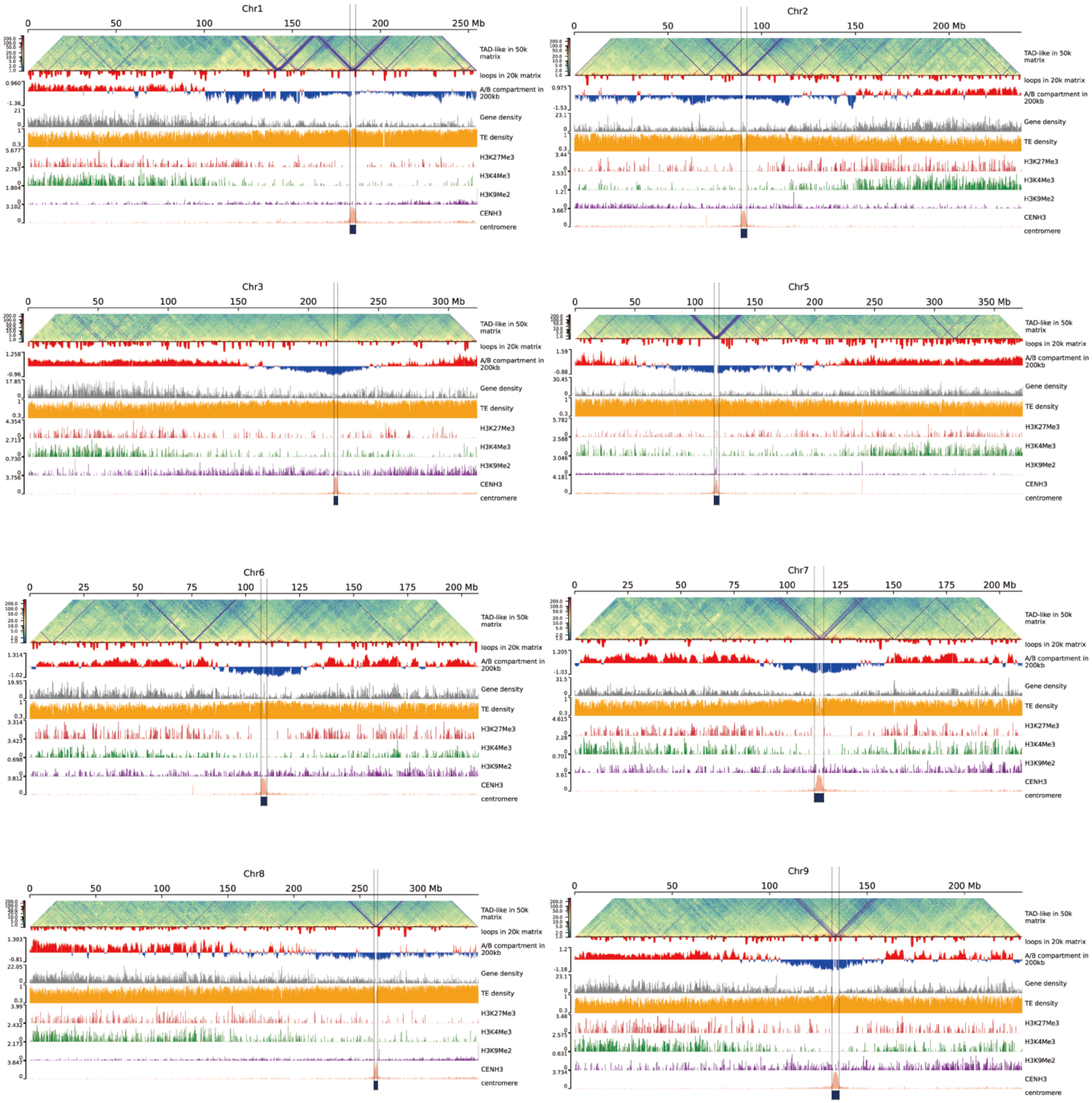


**Supplemental Figure 12. Epigenetic landscape of LsT2T genome.** From top to bottom of the plot, heatmap of TAD, density of 3D genomic loops and A/B compartments, gene density, TE density, the intensities of H3K27Me3, H3K4Me3, H3K9Me2 histone modifications, ChIP - seq marks and centromeric location were shown for each chromosome.

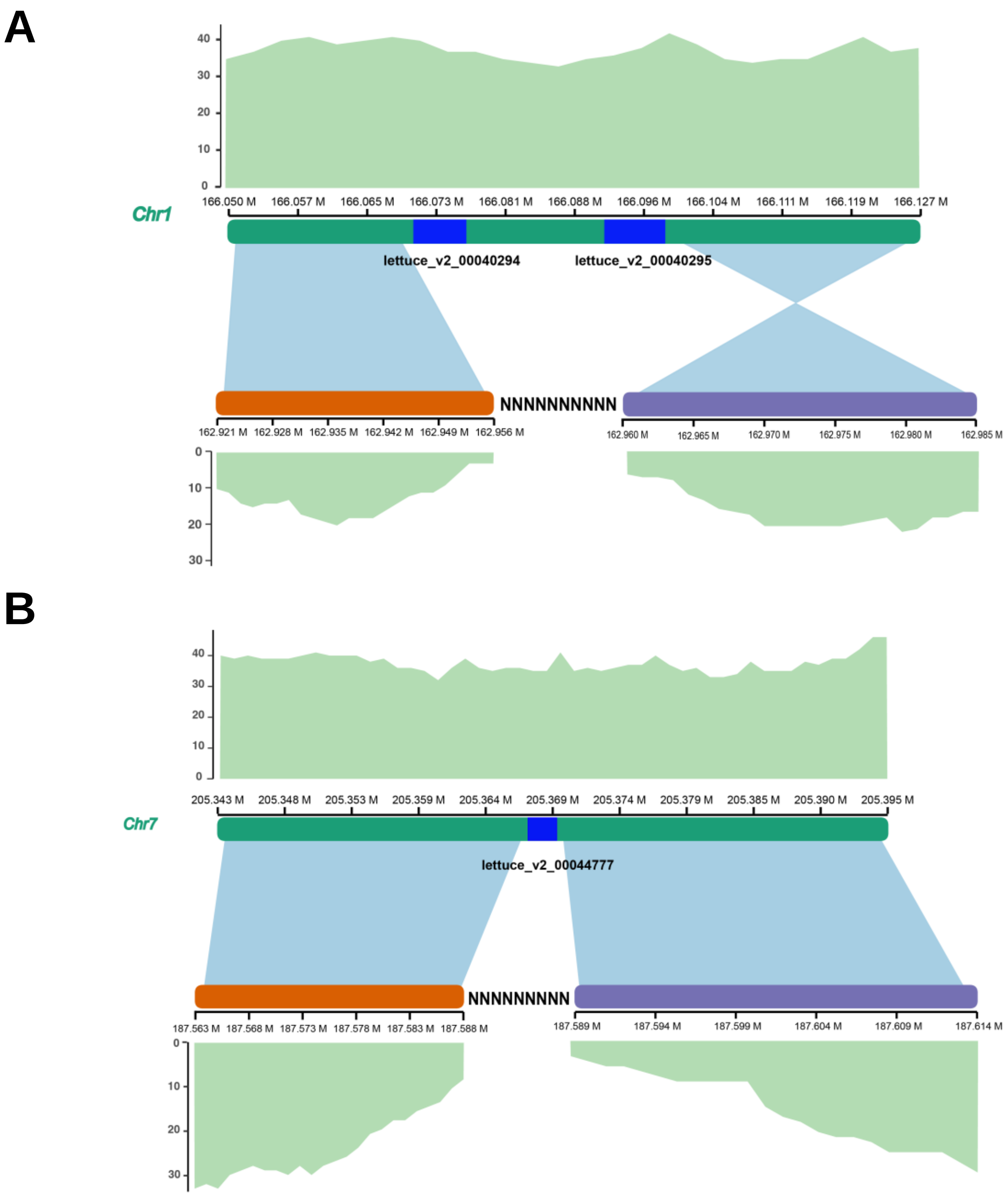


**Supplemental Figure 13. Diagram of three NLR genes identified in the gap filled regions of LsT2T genome.**

(**A-B**) Two genes on Chr1 **A** and one gene on Chr7 **B** were indicated by blue, respectively. The read coverages of ONT reads mapped to LsT2T (top) and *Salinas* genomes (bottom) were shown adjacent to the gaps and proximity regions.
