## Supplemental Table 1-8 for "The complete genome of lettuce reveals centromere landscape and empowers breeding"

**Supplementary Table 1. DNA sequencing data used for *de novo* genome assembly of *L. sativa* cv. *PKU06***

| | Total reads | Total bases | Coverage [a] | N50_len | Mapping rate |
|---|---|---|---|---|---|
| HiFi | 15,788,419 | 292,257,529,913 | 112.4 | 18,939 | 99.93% |
| ONT | 5,492,383 | 111,647,316,974 | 42.9 | 62,512 | 96.40% |
| NGS | 1,044,401,360 | 156,660,204,000 | 60.2 | 150 | 99.96% |
| Hi-C | 2,059,365,748 | 308,904,862,200 | 118.8 | 150 | 99.72% |

[a] The coverage depth (×) was calculated using the genome assembly size of 2.6 Gb for *L. sativa* .

**Supplementary Table 2. Comparison of genomic features of *Salinas* , *Angustana* and LsT2T genomes.**

| | *Salinas* | LsT2T |
|---|---|---|
| Total sequence length (bp) | 2,589,853,228 | 2,593,462,933 |
| Number of contig | 475 | 9 |
| Number of gaps | 384 | 0 |
| Contig N50 (Mb) | 12.5 | 283.1 |
| Longest contig (bp) | 410,295,809 | 409,415,085 |
| Number of telomeres assembled | 16 | 18 |
| Number of centromeres assembled | NA | 9 |
| Length of rDNA (NOR) assembled (Mb) | NA | 8.63 |
| Repeat content (%) | NA | 81.4 |
| Number of protein-coding genes | 44,232 | 45,507 |
| QV | 28.65 | 58.03 |
| BUSCO (%) | 98.4 | 97.6 |

NA: unavailable
QV: quality value
BUSCO: Benchmarking Universal Single-Copy Ortholog

**Supplementary Table 3. Coordinates of telomeres in LsT2T genome.**

| Chr | Start (bp) | End (bp) | Size (bp) | Start (bp) | End (bp) | Size (bp) |
|---|---|---|---|---|---|---|
| Chr1 | 1 | 7,531 | 7,530 | 258,115,905 | 258,130,948 | 15,043 |
| Chr2 | 1 | 77,665 | 77,664 | 238,524,979 | 238,535,665 | 10,686 |
| Chr3 | 1 | 3,638 | 3,637 | 320,634,812 | 320,657,391 | 22,579 |
| Chr4 | 1 | 24,626 | 24,625 | 409,325,439 | 409,415,085 | 89,646 |
| Chr5 | 1 | 44,211 | 44,210 | 373,630,944 | 373,650,124 | 19,180 |
| Chr6 | 1 | 23,108 | 23,107 | 207,787,258 | 207,794,748 | 7,490 |
| Chr7 | 1 | 16,889 | 16,888 | 210,585,715 | 210,595,814 | 10,099 |
| Chr8 | 1 | 8,254 | 8,253 | 345,091,913 | 345,107,305 | 15,392 |
| Chr9 | 1 | 6,025 | 6,024 | 229,566,750 | 229,575,853 | 9,103 |

**Supplementary Table 4. Coordinates  and composition of high-coveage region in LsT2T genome.**

| Chr | Start (bp) | End (bp) | Type | Coverage (200 kb) |
|---|---|---|---|---|
| Chr4 | 74,200,000.00 | 74,400,000.00 | chloroplast | 7,312.04 |
| Chr4 | 74,400,000.00 | 74,600,000.00 | chloroplast | 5,196.23 |
| Chr5 | 83,000,000.00 | 83,200,000.00 | chloroplast | 294.30 |
| Chr2 | 238,400,000.00 | 238,535,665.00 | rdna | 200.61 |
| Chr4 | 160,800,000.00 | 161,000,000.00 | rdna | 138.78 |
| Chr6 | 108,600,000.00 | 108,800,000.00 | mitochondria | 125.32 |
| Chr5 | 40,800,000.00 | 41,000,000.00 | mitochondria | 114.21 |
| Chr6 | 108,400,000.00 | 108,600,000.00 | mitochondria | 113.09 |
| Chr4 | 89,200,000.00 | 89,400,000.00 | chloroplast | 106.33 |
| Chr9 | 226,000,000.00 | 226,200,000.00 | mitochondria | 101.28 |
| Chr8 | 4,600,000.00 | 4,800,000.00 | chloroplast | 91.39 |
| Chr8 | 198,400,000.00 | 198,600,000.00 | mitochondria | 85.19 |
| Chr6 | 192,000,000.00 | 192,200,000.00 | mitochondria | 82.21 |

**Supplementary Table 5. Characterization of REs in LsT2T genome.**

| | *L. sativa* | |
|---|---|---|
| Type | Element length | Percentage |
| Retroelements | 1748503586 bp | 67.42% |
| SINEs | 12808171 bp | 0.49% |
| LINEs | 36472971 bp | 1.41% |
| L2/CR1/Rex | 2145928 bp | 0.08% |
| L1/CIN4 | 15180118 bp | 0.59% |
| CRE/SLACS | 2251700 bp | 0.09% |
| LTR elements | 1699222444 bp | 65.52% |
| Ty1/Copia | 706133068 bp | 27.23% |
| Gypsy/DIRS1 | 981316777 bp | 37.84% |
| Retroviral | 6191541 bp | 0.24% |
| DNA transposons: | 216358262 bp | 8.34% |
| hobo-Activator | 72808939 bp | 2.81% |
| Tc1-IS630-Pogo | 18906069 bp | 0.73% |
| Tourist/Harbinger | 31406854 bp | 1.21% |
| MULE-MUDR | 18747964 bp | 0.72% |
| Rolling-circles | 1793565 bp | 0.07% |
| Other (Mirage,P-element, Transib) | 11765309 bp | 0.45% |
| Unclassified: | 88189183 bp | 3.40% |
| Small RNA: | 189948 bp | 0.01% |
| Satellites: | 26006898 bp | 1.00% |
| Simple repeats: | 28762937 bp | 1.11% |
| Low complexity: | 2528432 bp | 0.10% |
| Total | 2112146175 bp | 81.44% |

**Supplementary Table 6. Annotaions of protein-encoding gene in LsT2T and v11 genomes.**

| | LsT2T | *Sallinas* |
|---|---|---|
| Number of protein coding genes | 45,507 | 44,232 |
| Number of unanchored genes | 0 | 485 |
| Average exon number | 4.386 | 5.218 |
| Average exon length (bp) | 248 | 260 |
| Average transcript length (bp) | 1,206 | 1,151 |
| Average gene length (bp) | 2,723 | 2,430 |
| Number of GO annotated genes | 16,464 | 16,464 |
| Number of KO annotated gene number | 16,929 | 16,389 |
| TPM > 1 | 26,084 | NA |

**Supplemental Table 7. Coordinate and composition of centromere in LsT2T genome.**

| Chr | Start (bp) | End (bp) | Size (bp) | L/S | DNA (%) | LINE (%) | Low complexity (%) | Copia (%) |
|---|---|---|---|---|---|---|---|---|
| Chr1 | 182699531 | 185911258 | 3211727 | 2.529775 | 3.30 | 2.08 | 0.06 | 6.26 |
| Chr2 | 88942479 | 92160659 | 3218180 | 1.645727 | 6.85 | 3.41 | 0.02 | 10.03 |
| Chr3 | 218237282 | 221018516 | 2781234 | 2.190282 | 8.09 | 2.53 | 0.06 | 30.87 |
| Chr4 | 274382309 | 277914143 | 3531834 | 2.088365 | 3.43 | 1.37 | 0.03 | 2.59 |
| Chr5 | 116080000 | 120300123 | 4220123 | 2.182547 | 4.76 | 2.84 | 0.05 | 7.39 |
| Chr6 | 107065176 | 109801206 | 2736030 | 1.092574 | 8.44 | 2.72 | 0.04 | 17.38 |
| Chr7 | 112782001 | 117265064 | 4483063 | 1.208412 | 6.97 | 2.54 | 0.03 | 13.24 |
| Chr8 | 260739975 | 263574706 | 2834731 | 3.197984 | 5.33 | 2.19 | 0.05 | 11.00 |
| Chr9 | 131940743 | 135756805 | 3816062 | 1.406332 | 8.30 | 3.28 | 0.04 | 13.41 |

| Gypsy (%) | Helitron (%) | Satellite (%) | Simple repeat (%) | SINE (%) | Unknown repeat (%) | Unknown (%) |
|---|---|---|---|---|---|---|
| 23.03 | 0.00 | 60.14 | 0.14 | 0.04 | 1.68 | 1.70 |
| 48.07 | 0.00 | 22.26 | 0.33 | 0.01 | 2.96 | 3.04 |
| 45.50 | 0.04 | 3.25 | 0.49 | 0.63 | 4.23 | 4.36 |
| 51.59 | 0.02 | 36.76 | 0.19 | 0.16 | 1.23 | 1.24 |
| 69.42 | 0.00 | 8.62 | 0.23 | 0.00 | 2.23 | 2.68 |
| 54.87 | 0.00 | 5.14 | 0.39 | 0.01 | 3.57 | 3.77 |
| 52.77 | 0.00 | 9.97 | 0.32 | 0.05 | 3.30 | 3.38 |
| 39.77 | 0.00 | 32.61 | 0.27 | 0.00 | 2.24 | 2.31 |
| 60.24 | 0.00 | 9.59 | 0.27 | 0.14 | 3.25 | 3.29 |

**Supplementary Table 8. 58 up-regulated NLR DEGs**

| Gene | TYPE | log2FC | Time | log2FC | Time | log2FC | Time | log2FC |
|---|---|---|---|---|---|---|---|---|
| lettuce_v2_00041910 | TIR-NBARC-LRR | 0.461610538 | 9hpi | 0.064702052 | 12hpi | 0.764311495 | 15hpi | 0.906896898 |
| lettuce_v2_00041380 | TIR-NBARC-LRR | - | 9hpi | - | 12hpi | - | 15hpi | - |
| lettuce_v2_00041350 | TIR-NBARC-LRR | 0.25452891 | 9hpi | -0.033030917 | 12hpi | 0.061680499 | 15hpi | 0.576663236 |
| lettuce_v2_00040881 | TIR-NBARC-LRR | 0.59826033 | 9hpi | -0.13166437 | 12hpi | 1.443785242 | 15hpi | 2.500815009 |
| lettuce_v2_00040878 | TIR-NBARC-LRR | 0.911316185 | 9hpi | -0.182713543 | 12hpi | 1.540734431 | 15hpi | 2.63248638 |
| lettuce_v2_00040875 | TIR-NBARC-LRR | 0.582937769 | 9hpi | -0.591290317 | 12hpi | 0.765425156 | 15hpi | 1.370005979 |
| lettuce_v2_00040874 | NBARC-LRR | 0.634702917 | 9hpi | -0.238546297 | 12hpi | 0.625673035 | 15hpi | 1.674789785 |
| lettuce_v2_00040808 | TIR-NBARC-LRR | 0.763391106 | 9hpi | 0.674886231 | 12hpi | 0.366641568 | 15hpi | 0.330091109 |
| lettuce_v2_00040790 | TIR-NBARC-LRR | 0.060548785 | 9hpi | -0.394115855 | 12hpi | 0.456709998 | 15hpi | 0.371213916 |
| lettuce_v2_00040783 | TIR-NBARC | 0.196060161 | 9hpi | 0.066270586 | 12hpi | 0.251312513 | 15hpi | 0.486809613 |
| lettuce_v2_00040764 | TIR-NBARC-LRR | 0.003480286 | 9hpi | -0.746661706 | 12hpi | 0.429803879 | 15hpi | - |
| lettuce_v2_00040762 | TIR-NBARC-LRR | 0.588320697 | 9hpi | -0.478385301 | 12hpi | 0.525367653 | 15hpi | 0.704036173 |
| lettuce_v2_00040760 | TIR-NBARC-LRR | 0.668103657 | 9hpi | -0.319733456 | 12hpi | 0.423282046 | 15hpi | 2.290743788 |
| lettuce_v2_00040758 | TIR-NBARC-LRR | 0.757552195 | 9hpi | -0.772273715 | 12hpi | 1.747959992 | 15hpi | 2.261114239 |
| lettuce_v2_00040750 | TIR-NBARC-LRR | 0.32291079 | 9hpi | 0.199802691 | 12hpi | -0.102253371 | 15hpi | 0.563369657 |
| lettuce_v2_00040617 | CC-NBARC-LRR | - | 9hpi | - | 12hpi | - | 15hpi | 2.414310312 |
| lettuce_v2_00040484 | TIR-NBARC-LRR | 0.428064699 | 9hpi | 0.196053322 | 12hpi | 0.775827165 | 15hpi | 1.155458722 |
| lettuce_v2_00040465 | TIR-NBARC-LRR | -0.203081461 | 9hpi | -0.331814808 | 12hpi | 0.586553205 | 15hpi | -0.449541815 |
| lettuce_v2_00040441 | TIR-NBARC-LRR | - | 9hpi | - | 12hpi | - | 15hpi | 4.154925825 |
| lettuce_v2_00040432 | TIR-NBARC-LRR | 1.312022172 | 9hpi | 1.397246685 | 12hpi | 2.191985983 | 15hpi | 4.078977393 |
| lettuce_v2_00040430 | TIR-NBARC-LRR | 1.303619594 | 9hpi | 0.62370251 | 12hpi | 0.680819643 | 15hpi | 0.731531654 |
| lettuce_v2_00039117 | TIR-NBARC-LRR | - | 9hpi | - | 12hpi | - | 15hpi | - |
| lettuce_v2_00039112 | TIR-NBARC | - | 9hpi | - | 12hpi | - | 15hpi | - |
| lettuce_v2_00035794 | TIR-NBARC-LRR | 1.690105721 | 9hpi | 1.396259824 | 12hpi | 1.275818045 | 15hpi | 1.997825573 |
| lettuce_v2_00001624 | TIR-NBARC-LRR | 1.501162526 | 9hpi | 0.925389123 | 12hpi | 2.10747378 | 15hpi | 2.296885119 |
| lettuce_v2_00006425 | TIR-NBARC-LRR | 0.002869563 | 9hpi | -0.069570574 | 12hpi | -0.20346129 | 15hpi | -0.344586287 |
| lettuce_v2_00006447 | TIR-NBARC-LRR | - | 9hpi | - | 12hpi | 5.721114727 | 15hpi | - |
| lettuce_v2_00051320 | CC-NBARC-LRR | 0.287670475 | 9hpi | -0.171662869 | 12hpi | 1.402908091 | 15hpi | 1.914037994 |
| lettuce_v2_00047635 | TIR-NBARC-LRR | - | 9hpi | - | 12hpi | - | 15hpi | - |
| lettuce_v2_00047579 | TIR-NBARC | 0.168618367 | 9hpi | -0.02258307 | 12hpi | 0.153494931 | 15hpi | 0.891734186 |
| lettuce_v2_00047568 | TIR-NBARC-LRR | 0.670210431 | 9hpi | -0.020967135 | 12hpi | 0.356628796 | 15hpi | 0.942292102 |
| lettuce_v2_00047567 | TIR-NBARC-LRR | 0.258863836 | 9hpi | -0.147359533 | 12hpi | 0.23921853 | 15hpi | 0.273643279 |

| | | | | | | | | |
|---|---|---|---|---|---|---|---|---|
| lettuce_v2_00047566 | TIR-NBARC-LRR | 0.269722626 | 9hpi | 0.063819305 | 12hpi | 0.702904161 | 15hpi | 1.386463247 |
| lettuce_v2_00047564 | TIR-NBARC | 0.194932562 | 9hpi | -0.130678138 | 12hpi | 0.535967504 | 15hpi | 0.207458113 |
| lettuce_v2_00044777 | TIR-NBARC | - | 9hpi | - | 12hpi | - | 15hpi | - |
| lettuce_v2_00016211 | TIR-NBARC-LRR | 0.669272922 | 9hpi | 0.31561276 | 12hpi | 0.900804685 | 15hpi | 1.224558747 |
| lettuce_v2_00016212 | TIR-NBARC-LRR | - | 9hpi | 0.977157055 | 12hpi | 2.184900736 | 15hpi | 2.296692421 |
| lettuce_v2_00017184 | TIR-NBARC-LRR | 0.756171806 | 9hpi | 0.441080617 | 12hpi | 2.199515217 | 15hpi | 3.045433668 |
| lettuce_v2_00017551 | TIR-NBARC | 0.268917505 | 9hpi | -0.671284447 | 12hpi | 1.308129101 | 15hpi | 1.653367263 |
| lettuce_v2_00017552 | TIR-NBARC | 0.752520921 | 9hpi | 0.16174918 | 12hpi | 1.461636193 | 15hpi | 2.355801537 |
| lettuce_v2_00017572 | TIR-NBARC-LRR | -0.463640604 | 9hpi | -0.041124007 | 12hpi | 0.85154442 | 15hpi | 0.406725101 |
| lettuce_v2_00017601 | TIR-NBARC-LRR | -0.122674889 | 9hpi | -0.460804592 | 12hpi | - | 15hpi | - |
| lettuce_v2_00017605 | TIR-NBARC | 0.750944537 | 9hpi | 0.095969523 | 12hpi | 1.325292226 | 15hpi | 3.752884885 |
| lettuce_v2_00017613 | TIR-NBARC-LRR | - | 9hpi | 0.008244845 | 12hpi | 1.31425585 | 15hpi | 1.562246938 |
| lettuce_v2_00017622 | TIR-NBARC | 0.598248207 | 9hpi | -0.176389207 | 12hpi | 0.074569631 | 15hpi | 0.536483699 |
| lettuce_v2_00020098 | TIR-NBARC-LRR | 0.540332756 | 9hpi | 0.003992047 | 12hpi | 0.161630867 | 15hpi | 0.344457497 |
| lettuce_v2_00020173 | TIR-NBARC-LRR | 2.801976975 | 9hpi | 4.581488227 | 12hpi | 4.59499701 | 15hpi | 6.200644484 |
| lettuce_v2_00020419 | CC-NBARC-LRR | -0.097127788 | 9hpi | 0.10028126 | 12hpi | 0.214667422 | 15hpi | 0.926306727 |
| lettuce_v2_00020420 | CC-NBARC-LRR | 0.442867419 | 9hpi | 0.239107224 | 12hpi | -0.207886833 | 15hpi | 1.388169868 |
| lettuce_v2_00020581 | CC-NBARC-LRR | - | 9hpi | 0.544513894 | 12hpi | 0.591517557 | 15hpi | 1.407747127 |
| lettuce_v2_00031868 | CC-NBARC-LRR | 0.940260337 | 9hpi | 1.284913665 | 12hpi | 2.010202094 | 15hpi | 3.152409708 |
| lettuce_v2_00031586 | NBARC-LRR | 0.113928568 | 9hpi | -0.190943597 | 12hpi | 0.229163833 | 15hpi | 0.548391235 |
| lettuce_v2_00030902 | TIR-NBARC | -0.266341915 | 9hpi | -0.252651952 | 12hpi | 0.920100446 | 15hpi | 0.688398656 |
| lettuce_v2_00030895 | TIR-NBARC | - | 9hpi | - | 12hpi | - | 15hpi | 2.325783474 |
| lettuce_v2_00029769 | TIR-NBARC-LRR | - | 9hpi | - | 12hpi | - | 15hpi | - |
| lettuce_v2_00029714 | TIR-NBARC-LRR | 1.067845432 | 9hpi | -0.104798735 | 12hpi | 1.146370091 | 15hpi | 1.355497828 |
| lettuce_v2_00029657 | TIR-NBARC | - | 9hpi | - | 12hpi | - | 15hpi | - |
| lettuce_v2_00029479 | TIR-NBARC-LRR | 0.082539023 | 9hpi | 0.372034556 | 12hpi | 1.940393019 | 15hpi | 1.742951828 |

| Time | log2FC | Time | log2FC | Time | log2FC | Time | log2FC | Time | log2FC | Time |
|---|---|---|---|---|---|---|---|---|---|---|
| 18hpi | 1.538196064 | 21hpi | 1.13264032 | 24hpi | 0.992575033 | 27hpi | 1.035396364 | 30hpi | 0.772576241 | 33hpi |
| 18hpi | 0.725048849 | 21hpi | 1.194674641 | 24hpi | - | 27hpi | 2.648637533 | 30hpi | 2.670726724 | 33hpi |
| 18hpi | 0.693938823 | 21hpi | 0.650800895 | 24hpi | 0.367380756 | 27hpi | 0.261415104 | 30hpi | 0.036188733 | 33hpi |
| 18hpi | 2.059550092 | 21hpi | 1.832314648 | 24hpi | 1.562962144 | 27hpi | 1.881388595 | 30hpi | 1.777529651 | 33hpi |
| 18hpi | 1.778063954 | 21hpi | 1.013154762 | 24hpi | 0.640125438 | 27hpi | 1.108929742 | 30hpi | 0.87913946 | 33hpi |
| 18hpi | 1.248829149 | 21hpi | 1.127712255 | 24hpi | 0.629253615 | 27hpi | 0.980618821 | 30hpi | 1.005686726 | 33hpi |
| 18hpi | 1.159532471 | 21hpi | 0.589693877 | 24hpi | 0.270449633 | 27hpi | 0.326475318 | 30hpi | 0.234369921 | 33hpi |
| 18hpi | 1.17628949 | 21hpi | 0.308644609 | 24hpi | 0.790570089 | 27hpi | 0.647414695 | 30hpi | 1.52834608 | 33hpi |
| 18hpi | 0.960087146 | 21hpi | 0.786299362 | 24hpi | 0.768929264 | 27hpi | 0.947889133 | 30hpi | 1.586355081 | 33hpi |
| 18hpi | 1.093329341 | 21hpi | 0.882841078 | 24hpi | 0.619362262 | 27hpi | 0.780385746 | 30hpi | 0.876287535 | 33hpi |
| 18hpi | - | 21hpi | - | 24hpi | 0.04462348 | 27hpi | 0.688667497 | 30hpi | -0.021347087 | 33hpi |
| 18hpi | 0.984934729 | 21hpi | 1.213525396 | 24hpi | 0.49158098 | 27hpi | 1.362378014 | 30hpi | 0.488653646 | 33hpi |
| 18hpi | 1.688297246 | 21hpi | 0.924930632 | 24hpi | 0.373675267 | 27hpi | 0.791693973 | 30hpi | -0.100973345 | 33hpi |
| 18hpi | 2.205864146 | 21hpi | 3.070663978 | 24hpi | 1.748194046 | 27hpi | 1.905852028 | 30hpi | 1.247688872 | 33hpi |
| 18hpi | 0.487609576 | 21hpi | 1.249399862 | 24hpi | 0.237195867 | 27hpi | 0.426865821 | 30hpi | -0.285438203 | 33hpi |
| 18hpi | 3.422881608 | 21hpi | 3.327990134 | 24hpi | 2.266277206 | 27hpi | 2.93825922 | 30hpi | 1.649061651 | 33hpi |
| 18hpi | 1.910834838 | 21hpi | 1.724148101 | 24hpi | 1.548942007 | 27hpi | 2.176474432 | 30hpi | 1.065604739 | 33hpi |
| 18hpi | 1.371309752 | 21hpi | 2.067820095 | 24hpi | 0.779633442 | 27hpi | 2.370198694 | 30hpi | -0.227872225 | 33hpi |
| 18hpi | 4.41618049 | 21hpi | 3.632202131 | 24hpi | 3.029007059 | 27hpi | 3.991683723 | 30hpi | 4.275566985 | 33hpi |
| 18hpi | 3.200469317 | 21hpi | 2.703512773 | 24hpi | 2.599633106 | 27hpi | 2.556683092 | 30hpi | 1.808388129 | 33hpi |
| 18hpi | 2.051455173 | 21hpi | 0.991094924 | 24hpi | 1.613497575 | 27hpi | 1.892560896 | 30hpi | 1.436079223 | 33hpi |
| 18hpi | - | 21hpi | 0.21445421 | 24hpi | - | 27hpi | 0.468213423 | 30hpi | 1.765733637 | 33hpi |
| 18hpi | 2.397412002 | 21hpi | 3.408771755 | 24hpi | 2.799553404 | 27hpi | 4.416587036 | 30hpi | 2.979582262 | 33hpi |
| 18hpi | 2.821378741 | 21hpi | 3.022244613 | 24hpi | 2.516123532 | 27hpi | 2.427971275 | 30hpi | 1.835883267 | 33hpi |
| 18hpi | 3.53477223 | 21hpi | 2.892039136 | 24hpi | 2.961884914 | 27hpi | 2.659462155 | 30hpi | 0.381941632 | 33hpi |
| 18hpi | 0.246550643 | 21hpi | 0.655808727 | 24hpi | 0.522309644 | 27hpi | 0.83607726 | 30hpi | 1.197407363 | 33hpi |
| 18hpi | 7.294609792 | 21hpi | 5.250508241 | 24hpi | 3.925441769 | 27hpi | 5.223863945 | 30hpi | 5.130750386 | 33hpi |
| 18hpi | 2.804423131 | 21hpi | 2.990240771 | 24hpi | 2.226267329 | 27hpi | 2.787103836 | 30hpi | 0.772411038 | 33hpi |
| 18hpi | - | 21hpi | 3.194023163 | 24hpi | 2.783018174 | 27hpi | - | 30hpi | - | 33hpi |
| 18hpi | 1.668741959 | 21hpi | 1.674121486 | 24hpi | 1.597879688 | 27hpi | 0.919972936 | 30hpi | 1.122465553 | 33hpi |
| 18hpi | 1.599891231 | 21hpi | 1.815768181 | 24hpi | 1.593498611 | 27hpi | 1.191637415 | 30hpi | 1.659244997 | 33hpi |
| 18hpi | 1.141883805 | 21hpi | 1.028071529 | 24hpi | 1.01440488 | 27hpi | 1.387100249 | 30hpi | 1.893664908 | 33hpi |

| | | | | | | | | | | |
|---|---|---|---|---|---|---|---|---|---|---|
| 18hpi | 1.791940832 | 21hpi | 1.540920883 | 24hpi | 1.101008085 | 27hpi | 1.18443611 | 30hpi | 0.390352562 | 33hpi |
| 18hpi | 1.34890043 | 21hpi | 1.196808089 | 24hpi | 1.129875345 | 27hpi | 1.269453115 | 30hpi | 1.405251446 | 33hpi |
| 18hpi | - | 21hpi | 5.270125552 | 24hpi | - | 27hpi | - | 30hpi | - | 33hpi |
| 18hpi | 3.034526174 | 21hpi | 2.993802013 | 24hpi | 2.061105515 | 27hpi | 2.345327233 | 30hpi | 2.348667617 | 33hpi |
| 18hpi | 4.76198267 | 21hpi | 4.886214258 | 24hpi | 3.790904651 | 27hpi | 3.702679622 | 30hpi | 3.525170529 | 33hpi |
| 18hpi | 4.409847789 | 21hpi | 3.88542839 | 24hpi | 3.185294213 | 27hpi | 3.599514225 | 30hpi | 2.972011123 | 33hpi |
| 18hpi | 1.099500969 | 21hpi | 1.51325013 | 24hpi | 0.943390571 | 27hpi | - | 30hpi | 0.198138293 | 33hpi |
| 18hpi | 2.898836005 | 21hpi | 2.75923836 | 24hpi | 2.590796242 | 27hpi | 2.357650617 | 30hpi | 1.850129596 | 33hpi |
| 18hpi | 1.348257295 | 21hpi | 1.326129685 | 24hpi | 1.332016131 | 27hpi | 1.251117979 | 30hpi | 1.110622417 | 33hpi |
| 18hpi | 1.008784569 | 21hpi | 1.49162526 | 24hpi | 0.135520753 | 27hpi | 0.242113113 | 30hpi | 0.294382709 | 33hpi |
| 18hpi | 2.229874476 | 21hpi | 2.09850074 | 24hpi | 1.524677076 | 27hpi | 1.263276575 | 30hpi | 0.837165051 | 33hpi |
| 18hpi | 3.220979916 | 21hpi | 2.686252695 | 24hpi | 2.631960919 | 27hpi | 2.821270238 | 30hpi | 3.595513402 | 33hpi |
| 18hpi | 1.51573502 | 21hpi | 1.719052189 | 24hpi | 1.439382311 | 27hpi | 1.407646325 | 30hpi | 1.971688767 | 33hpi |
| 18hpi | 0.846112029 | 21hpi | 0.997095306 | 24hpi | 0.996918512 | 27hpi | 1.18327229 | 30hpi | 1.571521864 | 33hpi |
| 18hpi | 7.689709346 | 21hpi | 5.822331469 | 24hpi | 5.644391878 | 27hpi | 5.309570293 | 30hpi | 4.050292335 | 33hpi |
| 18hpi | 2.368532712 | 21hpi | 1.786650803 | 24hpi | 1.079902939 | 27hpi | 1.320140767 | 30hpi | 1.136797013 | 33hpi |
| 18hpi | 1.900175877 | 21hpi | 2.112567148 | 24hpi | 1.104679629 | 27hpi | 1.409977575 | 30hpi | 2.232573766 | 33hpi |
| 18hpi | 3.603737375 | 21hpi | 2.740385303 | 24hpi | 4.228148794 | 27hpi | 3.958587913 | 30hpi | 2.933867156 | 33hpi |
| 18hpi | 4.082118778 | 21hpi | 3.96363636 | 24hpi | 2.839654727 | 27hpi | 2.887269302 | 30hpi | 2.48016554 | 33hpi |
| 18hpi | 1.195495716 | 21hpi | 1.077323022 | 24hpi | 0.733981024 | 27hpi | 1.063694341 | 30hpi | 0.991894081 | 33hpi |
| 18hpi | 1.192303645 | 21hpi | 1.343571794 | 24hpi | 0.503579493 | 27hpi | 0.989539566 | 30hpi | 0.404544841 | 33hpi |
| 18hpi | 2.464109434 | 21hpi | 1.751812446 | 24hpi | 0.66539446 | 27hpi | 1.654760202 | 30hpi | 0.895000385 | 33hpi |
| 18hpi | - | 21hpi | 7.824809954 | 24hpi | 7.113393984 | 27hpi | 8.081047154 | 30hpi | 7.298363634 | 33hpi |
| 18hpi | 2.545430559 | 21hpi | 2.527155378 | 24hpi | 1.361865798 | 27hpi | 2.260006236 | 30hpi | 1.855272879 | 33hpi |
| 18hpi | 2.727335798 | 21hpi | - | 24hpi | 1.317365557 | 27hpi | - | 30hpi | 0.738750361 | 33hpi |
| 18hpi | 3.085183474 | 21hpi | 3.055989977 | 24hpi | 2.363331033 | 27hpi | 2.206228136 | 30hpi | 3.374085133 | 33hpi |

| log2FC | Time | log2FC | Time | log2FC | Time | log2FC | Time | log2FC | Time |
|---|---|---|---|---|---|---|---|---|---|
| 0.678166531 | 36hpi | 1.011796508 | 39hpi | 1.130194353 | 42hpi | 0.288189343 | 45hpi | 0.238233728 | 48hpi |
| 2.343411886 | 36hpi | 3.226004531 | 39hpi | 2.65543294 | 42hpi | 2.435703356 | 45hpi | 1.919503735 | 48hpi |
| -0.023134041 | 36hpi | 0.498664351 | 39hpi | 1.017928979 | 42hpi | -0.101512554 | 45hpi | -0.2897194 | 48hpi |
| 1.434089471 | 36hpi | 1.547847509 | 39hpi | 1.265007313 | 42hpi | 0.657852076 | 45hpi | 0.324966383 | 48hpi |
| 0.646542396 | 36hpi | 1.395863231 | 39hpi | 0.764492099 | 42hpi | -0.152006632 | 45hpi | 0.005535867 | 48hpi |
| 0.398748393 | 36hpi | 1.098304499 | 39hpi | 0.782057282 | 42hpi | 0.18131688 | 45hpi | 0.295385288 | 48hpi |
| 0.24262091 | 36hpi | 0.68299062 | 39hpi | 0.184973905 | 42hpi | -0.786921472 | 45hpi | -0.427417564 | 48hpi |
| 1.225854004 | 36hpi | 1.707852016 | 39hpi | 1.139786598 | 42hpi | 0.817526543 | 45hpi | 0.33564853 | 48hpi |
| 0.753865874 | 36hpi | 1.140191415 | 39hpi | 1.029472427 | 42hpi | 0.582419368 | 45hpi | 0.536275075 | 48hpi |
| 0.326810436 | 36hpi | 0.646484511 | 39hpi | 0.82360204 | 42hpi | -0.059524119 | 45hpi | -0.144325943 | 48hpi |
| 0.457915943 | 36hpi | 0.731865303 | 39hpi | 2.048514043 | 42hpi | 0.082527506 | 45hpi | -0.128706446 | 48hpi |
| -0.353810002 | 36hpi | 0.137340292 | 39hpi | -0.860306765 | 42hpi | 0.508395437 | 45hpi | 0.269033698 | 48hpi |
| -0.898307392 | 36hpi | -0.267616432 | 39hpi | -0.470429072 | 42hpi | -0.768883776 | 45hpi | -0.746344788 | 48hpi |
| 1.489146684 | 36hpi | 1.956879962 | 39hpi | 1.979043543 | 42hpi | 0.676586841 | 45hpi | 0.445528235 | 48hpi |
| 0.005127389 | 36hpi | 0.145741006 | 39hpi | 0.215580425 | 42hpi | -0.240686912 | 45hpi | -0.620657844 | 48hpi |
| 1.984114825 | 36hpi | 3.213523227 | 39hpi | 2.028324671 | 42hpi | 2.080936655 | 45hpi | 1.797564897 | 48hpi |
| 1.348014687 | 36hpi | 1.475685043 | 39hpi | 1.298019254 | 42hpi | 0.938977955 | 45hpi | 0.582121104 | 48hpi |
| -0.042810032 | 36hpi | -0.094590995 | 39hpi | 1.71096527 | 42hpi | 1.036271918 | 45hpi | 0.889588608 | 48hpi |
| 2.444135752 | 36hpi | 4.989753468 | 39hpi | 2.886834546 | 42hpi | 4.264284974 | 45hpi | 3.506273207 | 48hpi |
| 1.438283583 | 36hpi | 1.865085588 | 39hpi | 2.913594879 | 42hpi | 0.974376964 | 45hpi | 2.107897618 | 48hpi |
| 0.713044042 | 36hpi | 2.10733226 | 39hpi | 1.42339131 | 42hpi | 0.918812582 | 45hpi | 1.462038182 | 48hpi |
| 2.68227503 | 36hpi | - | 39hpi | 1.183457859 | 42hpi | 1.477219256 | 45hpi | 0.652502977 | 48hpi |
| 2.775871735 | 36hpi | 3.241033296 | 39hpi | 1.930752135 | 42hpi | 2.418507445 | 45hpi | 1.498285152 | 48hpi |
| 1.681449708 | 36hpi | 2.152223744 | 39hpi | 1.625571634 | 42hpi | -0.090423171 | 45hpi | 0.255824328 | 48hpi |
| 0.823269532 | 36hpi | 1.028970315 | 39hpi | 1.098031548 | 42hpi | 0.478305114 | 45hpi | 1.098064189 | 48hpi |
| 0.858521768 | 36hpi | 1.531037455 | 39hpi | 0.652099263 | 42hpi | 0.773943899 | 45hpi | 0.212703405 | 48hpi |
| 5.895201119 | 36hpi | 5.404689996 | 39hpi | 3.671792883 | 42hpi | 3.210363076 | 45hpi | 3.321487601 | 48hpi |
| 0.530384993 | 36hpi | 1.117216916 | 39hpi | 0.717901102 | 42hpi | 0.27617031 | 45hpi | 0.734635944 | 48hpi |
| - | 36hpi | - | 39hpi | - | 42hpi | - | 45hpi | - | 48hpi |
| 0.832177384 | 36hpi | 1.224272417 | 39hpi | 0.778800445 | 42hpi | 0.10902622 | 45hpi | 0.736688463 | 48hpi |
| 0.692676449 | 36hpi | 1.381115711 | 39hpi | 0.839825284 | 42hpi | 0.728894234 | 45hpi | 0.712005577 | 48hpi |
| 1.02218256 | 36hpi | 1.402528785 | 39hpi | 1.006135616 | 42hpi | 0.789856222 | 45hpi | 0.567487216 | 48hpi |

| | | | | | | | | | |
|---|---|---|---|---|---|---|---|---|---|
| 0.219556776 | 36hpi | 0.544667624 | 39hpi | 0.292370435 | 42hpi | 0.071238265 | 45hpi | 0.187026798 | 48hpi |
| 0.693576082 | 36hpi | 1.129329536 | 39hpi | 0.646539839 | 42hpi | 0.230803492 | 45hpi | 0.221146801 | 48hpi |
| - | 36hpi | - | 39hpi | - | 42hpi | - | 45hpi | - | 48hpi |
| 1.589412138 | 36hpi | 1.976055346 | 39hpi | 2.103517242 | 42hpi | 1.8973738 | 45hpi | 1.584286819 | 48hpi |
| 2.196835614 | 36hpi | 3.024313143 | 39hpi | 1.497818595 | 42hpi | 2.052209765 | 45hpi | 2.608198748 | 48hpi |
| 2.012128491 | 36hpi | 2.63759892 | 39hpi | 2.833187313 | 42hpi | 2.32441583 | 45hpi | 2.258541973 | 48hpi |
| - | 36hpi | 0.781268846 | 39hpi | -1.209467193 | 42hpi | 1.116751608 | 45hpi | 0.376190621 | 48hpi |
| 1.932264417 | 36hpi | 2.38856091 | 39hpi | 1.128449454 | 42hpi | 0.906541231 | 45hpi | 1.15036538 | 48hpi |
| 1.160910866 | 36hpi | 1.40699389 | 39hpi | 1.230038258 | 42hpi | 0.54129291 | 45hpi | 0.544643116 | 48hpi |
| - | 36hpi | - | 39hpi | -0.647322078 | 42hpi | -0.050724272 | 45hpi | -0.167037078 | 48hpi |
| 1.710543961 | 36hpi | 2.326709756 | 39hpi | 1.451120669 | 42hpi | 0.478455325 | 45hpi | 1.162395923 | 48hpi |
| 1.869303943 | 36hpi | 3.246550778 | 39hpi | 2.166908586 | 42hpi | 2.21743776 | 45hpi | 1.756626956 | 48hpi |
| 1.187665842 | 36hpi | 1.457016684 | 39hpi | 1.228933716 | 42hpi | 0.554879747 | 45hpi | 0.623270673 | 48hpi |
| 0.79617817 | 36hpi | 1.18710478 | 39hpi | 0.688162897 | 42hpi | 0.291847393 | 45hpi | 0.30446343 | 48hpi |
| 3.949143133 | 36hpi | 4.27947483 | 39hpi | 3.772873769 | 42hpi | 1.954143281 | 45hpi | 4.164682784 | 48hpi |
| 0.964012325 | 36hpi | 1.218636745 | 39hpi | 0.445161117 | 42hpi | 0.629822497 | 45hpi | 0.468140691 | 48hpi |
| 0.62624839 | 36hpi | 1.361878605 | 39hpi | 0.754270044 | 42hpi | 0.675796721 | 45hpi | 0.819741234 | 48hpi |
| 3.826029868 | 36hpi | 4.302750069 | 39hpi | 3.159984296 | 42hpi | 2.90851971 | 45hpi | 3.32687018 | 48hpi |
| 2.160395965 | 36hpi | 2.346489198 | 39hpi | 1.969860285 | 42hpi | 0.573604745 | 45hpi | 1.301150455 | 48hpi |
| 0.350994432 | 36hpi | 0.385074317 | 39hpi | 0.134173302 | 42hpi | -0.006910087 | 45hpi | 0.059395781 | 48hpi |
| 0.254998343 | 36hpi | 0.301504821 | 39hpi | -0.175564989 | 42hpi | 0.612806175 | 45hpi | 0.039510149 | 48hpi |
| - | 36hpi | - | 39hpi | 1.079529033 | 42hpi | 1.709318387 | 45hpi | 0.952408653 | 48hpi |
| 6.52241778 | 36hpi | 7.679945787 | 39hpi | 6.836076432 | 42hpi | 7.203748999 | 45hpi | 7.570251162 | 48hpi |
| 1.935413956 | 36hpi | 2.254156085 | 39hpi | 1.724398912 | 42hpi | 1.463909881 | 45hpi | 1.920951314 | 48hpi |
| - | 36hpi | - | 39hpi | - | 42hpi | - | 45hpi | - | 48hpi |
| 1.791579836 | 36hpi | 2.87349141 | 39hpi | 2.357659347 | 42hpi | 1.53376853 | 45hpi | 1.269203214 | 48hpi |
